# Resting state global brain activity induces bias in fMRI motion estimates

**DOI:** 10.1101/2023.10.31.565023

**Authors:** Yixiang Mao, Conan Chen, Truong Nguyen, Thomas T. Liu

## Abstract

Head motion is a significant source of artifacts in resting-state fMRI (rsfMRI) studies and has been shown to affect resting-state functional connectivity (rsFC) measurements. In many rsfMRI studies, motion parameters estimated from volume registration are used to characterize head motion and to mitigate motion artifacts in rsfMRI data. While prior task-based fMRI studies have shown that task-evoked brain activations may induce temporally correlated bias in the motion estimates, resulting in artificial activations after registration, relatively little is known about neural-related bias in rsfMRI motion parameters. In this study, we demonstrate that neural-related bias exists in rsfMRI motion estimates and characterize the potential effects of the bias on rsFC estimates. Using a public multi-echo rsfMRI dataset, we use the differences between motion estimates from the first echo and second echo data as a measure of neural-induced bias. We show that the resting-state global activity of the brain, as characterized with the global signal (GS), induces bias in the motion estimates in the y- and z-translational axes. Furthermore, we demonstrate that the GS-related bias reflects superior-inferior and anterior-posterior asymmetries in the GS beta coefficient map. Finally, we demonstrate that regression with biased motion estimates can negatively bias rsFC estimates and also reduce rsFC differences between young and old subjects.

## 1. Introduction

Head motion is a major source of artifacts in fMRI data, and motion estimates from data registration are commonly used to characterize and mitigate motion-related artifacts (Friston et al., 1996; Power et al., 2012; Satterthwaite et al., 2012; Dijk et al., 2012; Yan et al., 2013; Power et al., 2014, 2015). Prior task-based fMRI studies have demonstrated that task-evoked brain activations can induce bias in motion estimates that is temporally correlated with brain activations (Freire and Mangin, 2001; Freire et al., 2002). In contrast, relatively little is known about neural-related bias in motion estimates from resting-state fMRI (rsfMRI) data.

Detecting such bias is challenging due to the absence of ground truth head motion and neural signals in rsfMRI studies. To first order, fMRI signal changes resulting from head motion and neural activity can be distinguished based on their dependence on echo time (TE): neural activity causes TE-dependent blood-oxygen-level-dependent (BOLD) fluctuations, whereas head motion largely contributes to TE-independent non-BOLD signal changes. Taking advantage of these differences, prior studies have demonstrated that BOLD and non-BOLD signals can be distinguished using multi-echo fMRI (MEfMRI) that acquires data at different TEs (Buur et al., 2009; Bright and Murphy, 2013; Kundu et al., 2012, 2017). For example, Burr et al. effectively used the first echo data acquired at a short TE with minimal BOLD-weighting to model and correct for motion artifacts in the BOLD-weighted second echo data (Buur et al., 2009).

Burr’s findings imply that it may be feasible to identify neural-related bias by comparing motion estimates obtained from the first and second echo data. Since head motion primarily results in TE-independent signal changes, its effects should be captured in the motion estimates derived from both the first and second echo data. In contrast, there is typically minimal BOLD weighting in the first echo data but strong BOLD weighting in the second echo data. As prior work has shown that a higher level of brain activation can lead to greater bias in motion estimates (Freire and Mangin, 2001; Freire et al., 2002), potential neural-related bias should exhibit greater magnitude in the motion estimates from the second echo data as compared to those from the first echo data. Taken together, potential neural-related bias may be isolated from head motion by examining the difference between the motion estimates obtained from the first and second echo data.

Prior task-based fMRI studies have shown that brain activations with a larger spatial extent can lead to a higher level of bias in the motion estimates (Freire and Mangin, 2001; Freire et al., 2002). This finding motivates us to examine whether global brain activity in rsfMRI leads to bias in the motion estimates. In this study, we used the global signal (GS), calculated as the mean fMRI signal over the voxels within the brain, as a proxy for global brain activity. While the interpretation of the GS is still controversial (reviewed in (Liu et al., 2017)), there is growing evidence suggesting that the GS is linked to global neural activity (Wong et al., 2013, 2016; Falahpour et al., 2016, 2018; Liu et al., 2018; Gu et al., 2021).

In this work, we characterized the bias in rsfMRI motion parameters as estimated by AFNI *3dvolreg*. Furthermore, we investigated the consequences of using biased estimates as regressors in resting-state functional connectivity (rsFC) analyses.

## 2. Methods

### 2.1. Subjects and MEfMRI data acquisition

We used a public dataset (denoted as the Cornell-York dataset and described in (Setton et al., 2022; Spreng et al., 2022)) downloaded from OpenNeuro (dataset ds003592). The Cornell-York dataset includes multi-echo fMRI data collected from 301 healthy subjects (181 younger and 120 older adults). The data from 238 subjects were acquired on a 3T GE Discovery MR750 MRI scanner with a 32-channel head coil. The data from the remaining 63 subjects were collected on a 3T Siemens Trio MRI scanner with a 32-channel head coil. For each subject, two 10-min resting-state runs were acquired using an ME EPI sequence on the GE scanner (204 volumes; TR=3000 ms; TE=13.7, 30, 47 ms; flip angle=83°; FOV=210 mm; voxel size=3 × 3 × 3 mm^3^; matrix size=72 × 72 × 46; SENSE acceleration factor = 2.5; phase encoding direction: A-P) or on the Siemens scanner (200 volumes; TR=3000 ms; TE=14, 29.96, 45.92 ms; flip angle=83°; FOV=216 mm; voxel size=3.4 × 3.4 × 3 mm^3^; matrix size=64 × 64 × 43; GRAPPA acceleration factor = 3; phase encoding direction: A-P). The subjects were instructed to stay awake and lie still with their eyes open during the scans.

### 2.2. Data preprocessing

AFNI was used for data preprocessing (Cox, 1996). The fMRI data from the first and second echoes were used and denoted as e1 and e2, respectively. The data were first reoriented to Right-Anterior-Inferior (RAI) orientation (AFNI *3dresample -orient RAI*), and the first 6 TRs of the data were discarded to allow magnetization to reach a steady state. For each run and each echo, the data were normalized so that the mean signal over all voxels and all volumes was equal to 100. Since the normalization is equivalent to multiplication by a global scaling factor, the spatial and temporal information of the data in each run is preserved.

### 2.3. Calculation of the motion estimates and global signal

In this study, we examined the motion estimates from AFNI *3dvolreg* (Cox and Jesmanowicz, 1999) using the first volume as the reference volume. The default weights used in registration were disabled by feeding an all-ones image to the `-weight` option. This approach weights all voxels equally during registration and simplifies the theory. As shown in Supplementary Material (Fig. S3), similar results were obtained when using the default weights. For each run, the motion was estimated separately for the e1 and e2 data, resulting in two sets of estimates. The motion parameters were multiplied by −1 to represent the movement of the volumes as compared to the reference volume (see Appendix A).

For each run, the global signal (GS) was calculated from the unregistered e2 data. As shown in Supplementary Material (Fig. S3), nearly identical results were obtained when the GS was calculated from the registered e2 data. Before calculating the GS, each voxel’s signal was converted to a percent change signal. Then, the GS was computed by averaging over all voxels within the brain (brain masks were formed by AFNI *3dAutomask*). Finally, the mean and the linear and quadratic trends were regressed out from the motion estimates and GS.

### 2.4. Identifying BOLD-weighted GS bias in the motion estimates

To characterize BOLD-weighted GS bias in the motion estimates, we considered a simple signal model for the e1 and e2 motion estimates. Let ***m***_*e*1_ ∈ ℝ^*K*×1^ and ***m***_*e*2_ ∈ ℝ^*K*×1^ represent the motion estimates of one motion axis each from e1 and e2, respectively, where *K* is the number of volumes. We model the motion estimates as the weighted sum of head motion, BOLD-weighted GS bias and estimation error,

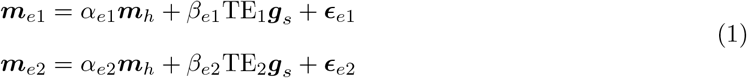

where ***m***_*h*_ ∈ ℝ^*K*×1^ represents head motion, *α*_*ei*_ is the regression weight corresponding to head motion for the *i*th echo, ***g***_*s*_ ∈ ℝ^*K*×1^ is the GS, TE_1_ and TE_2_ are the first and second echo times, respectively, *β*_*ei*_ is the regression weight of the BOLD-weighted GS bias for the *i*th echo and ***ϵ***_*ei*_ ∈ ℝ^*K*×1^ is the estimation error for the *i*th echo. With the above model, subtracting ***m***_*e*1_ from ***m***_*e*2_ yields

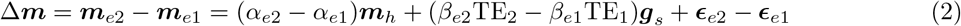

Since prior findings suggest that head motion can be accurately estimated from both e1 and e2 data (Buur et al., 2009; Speck and Hennig, 2001), we assume that *α*_*e*1_ ≈ *α*_*e*2_, yielding

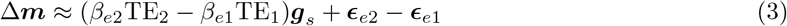

Note that Δ***m*** isolates BOLD-weighted GS bias from head motion. Therefore, we can examine the presence of potential bias by assessing the significance of the correlation between Δ***m*** and the GS. For each run and motion axis, we calculated the correlation between Δ***m*** and the GS (denoted as *r*(Δ***m***, GS)) and assessed the statistical significance of *r*(Δ***m***, GS) values on a per-run basis using empirical null distributions. For each motion axis, a null distribution was formed by calculating *r*(Δ***m***, GS) values using all possible permutations across runs, i.e. pairing the GS from one run to Δ***m*** from other runs and looping over all runs. The resulting null distributions consisted of 361,802 samples. Then, for each motion axis, we used the null distribution to compute the two-sided p-value associated with the *r*(Δ***m***, GS) value calculated from each run’s measured data. A p-value threshold of 0.05 was divided by 602 (the number of runs) to correct for multiple comparisons, and the Bonferroni corrected threshold was used to determine the significant correlations. Finally, we calculated the percentage of runs that show significant *r*(Δ***m***, GS) values for each motion axis.

In this study, we used the GS to represent the global activity of the brain. To reduce the potential effect of motion artifacts in the GS, we repeated the above analysis after regressing out the e1 motion regressors from both the GS and Δ***m***. The e1 motion regressors included the six motion parameters estimated from the e1 data and their first derivatives.

### 2.5. Spatial maps underlying r(Δ***m***, *GS*)

To provide insight into the mechanisms underlying BOLD-weighted GS bias, we derived an empirical approximation for *r*(Δ***m***, GS). As shown in Appendix C,

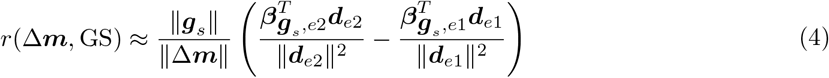

where 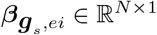 is the GS beta coefficient map for the *i*th echo, ***d***_*ei*_ ∈ ℝ^*N* ×1^ is the spatial derivative image with respect to (w.r.t.) one motion axis of the *i*th echo, *N* is the number of voxels and ∥ · ∥ denotes the *L*2-norm. For each run, the GS beta coefficient map 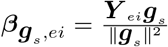 was calculated from the linear fit of the GS to the unregistered functional data ***Y*** _*ei*_ ∈ ℝ^*N* ×*K*^ of the *i*th echo.

We calculated the spatial derivative images w.r.t. the motion axes following the algorithm implemented in AFNI *3dvolreg*. For each run and each echo, the spatial derivative images were calculated based on the reference volume used in motion estimation. Denoting the reference image of the *i*th echo as ***y***_*r,ei*_ ∈ ℝ^*N* ×1^, the spatial derivative image of the *i*th echo w.r.t the *j*th motion axis was calculated as

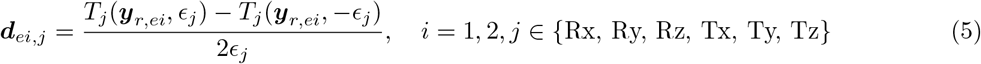

where *T*_*j*_: ℝ^*N* ×1^ → ℝ^*N* ×1^ is the function that transforms the reference volume along the *j*th motion axis, *ϵ*_*j*_ is the transformation parameter, Rx, Ry and Rz represent x-, y- and z-rotation, respectively and Tx, Ty, Tz represent x-, y- and z-translation, respectively. When calculating the spatial derivative images, the transformation was performed with AFNI *3drotate-heptic*. For the translational motion axes, *ϵ* Was set to 2.1 mm. For the rotational motion axes, *ϵ* was set to 0.4°. These *ϵ* values were determined by AFNI *3dvolreg* based on the spatial resolution of the functional data.

### 2.6. Effect of the bias on ROI-ROI FC via motion regression

In this work, we investigated the effect of regression with biased motion estimates on rsFC estimates between nodes within the default mode network (DMN) and task positive network (TPN). The regions of interest (ROIs) within these networks were defined in (Dijk et al., 2010), with four ROIs in the DMN (posterior cingulate cortex (PCC), lateral parietal cortex (LatPar), medial prefrontal cortex (mPFC) and Hippocampal formation (HF)) and three ROIs in the TPN (frontal eye field (FEF), intraparietal cortex (IPS) and middle temporal area (MT+)). Seed ROIs were created using a sphere with a radius of 12 mm centered about each seed coordinate. The left and right ROIs were combined to form bilateral ROIs. Prior to averaging signals within each ROI, the e2 data after volume registration were transferred to MNI space and spatially smoothed with a 4mm FWHM Gaussian kernel.

To reduce the confounding effects of head motion, motion censoring was performed before motion regression. For each run, framewise displacement (FD) was calculated based on six motion parameters estimated from the e1 data (the calculation of FD follows the description in (Power et al., 2012)). Volumes with FD values larger than 0.2 mm were censored. We show in Supplementary Material (Fig. S5 and S6) that motion censoring has little impact on the effect of regression with biased motion estimates on rsFC. After motion censoring, we calculated and compared the ROI-ROI FC after e2 and e1 motion regression to assess the effect of potential bias on FC analysis. For each ROI, the ROI-based seed signal was calculated by averaging the percent change BOLD signal over the voxels in that ROI. Then, the motion regressors, including the six motion parameters and their first derivatives were regressed out from the ROI signals. For each pair of ROIs, the ROI-ROI FC was computed as the Pearson’s correlation coefficient between the ROI average signals. Correlation values were converted to z-scores using the Fisher-z transformation. The differences in the ROI-ROI FC calculated after e2 and e1 motion regression were calculated to assess the effect of the bias on FC. The e2-e1 differences in r-values and z-scores were denoted as Δ*r* and Δ*z*, respectively.

Furthermore, we examined the effect of the bias on FC within four groups of runs based on levels of head motion and GS. For each run, the level of head motion was measured by the mean FD value calculated by averaging over the FD values across time. The level of the GS was measured by the GS amplitude (aGS) computed as the standard deviation of the GS after motion censoring. All the runs were first divided into two groups based on their mean FD values. Runs with mean FD values larger than the group median were classified as high motion runs and the remaining runs were classified as low motion runs. Then, within each FD-based group, the runs were further divided into two groups based on aGS. Runs with aGS values larger than the group median were classified as high aGS runs and the remaining runs were classified as low aGS runs. Consequently, we formed four groups of runs: 1) low motion and high aGS runs, (2) low motion and low aGS runs, (3) high motion and high aGS runs and (4) high motion and low aGS runs. A one-way ANOVA was calculated on mean Δ*z* over all ROI pairs to assess whether there was a group effect. Post-hoc two-sample t-tests were calculated to characterize the differences between pairs of groups.

Additionally, to verify that the effect of the bias on FC was dominated by the motion estimates from the axes where we found GS-induced bias, we evaluated the effect of two subsets of motion regressors. One of the subsets included the motion estimates and their first derivatives from the Ty and Tz axes (where we found GS-induced bias), while the other subset included the motion regressors from the other four motion axes, where minimal GS-induced was observed.

### 2.7. Effect of the bias on young vs. old group-level FC analysis

We first examined whether the bias in motion estimates affects the ROI-ROI FC of the young and old subjects differently by comparing Δ*r* and Δ*z* between the young and old runs. Furthermore, we investigated if the bias alters the group-level FC analysis between the young and old subjects. For each pair of ROIs, the significance of the FC differences between the young and old subjects was assessed by a permutation test with 1 × 10^7^ random permutations to allow us to apply p-value thresholds of 0.01, 1 × 10^−3^ and 1 × 10^−6^. Also, the effect size of the differences was measured with Cohen’s *d*. The FC differences and the significance and effect size of the differences calculated after e1 and e2 motion regression were compared.

## 3. Results

### 3.1. Examples of the GS and motion estimates

Fig. 1 (a,b) show the motion estimates in the Tz axis, including ***m***_*e*1_ (blue), ***m***_*e*2_ (green) and Δ***m*** (red) from two example runs with (a) low and (b) high levels of motion. Note that for the high motion run, ***m***_*e*1_ and ***m***_*e*2_ are plotted at 1*/*20th scale to facilitate comparison with ***m***_*e*1_ and ***m***_*e*2_ from the low motion run. As shown in these subfigures, Δ***m*** estimates from both runs fluctuate in a similar range from −0.05 to 0.05 mm (std(Δ***m***) = 0.017 and 0.025 for the low and high motion runs, respectively). In contrast, the standard deviations of ***m***_*e*1_ and ***m***_*e*2_ from the high motion run (std(***m***_*e*1_) = 0.583, std(***m***_*e*2_) = 0.602) are an order of magnitude larger than the standard deviations of ***m***_*e*1_ and ***m***_*e*2_ from the low motion run (std(***m***_*e*1_) = 0.027, std(***m***_*e*2_) = 0.034).

**Figure 1:**
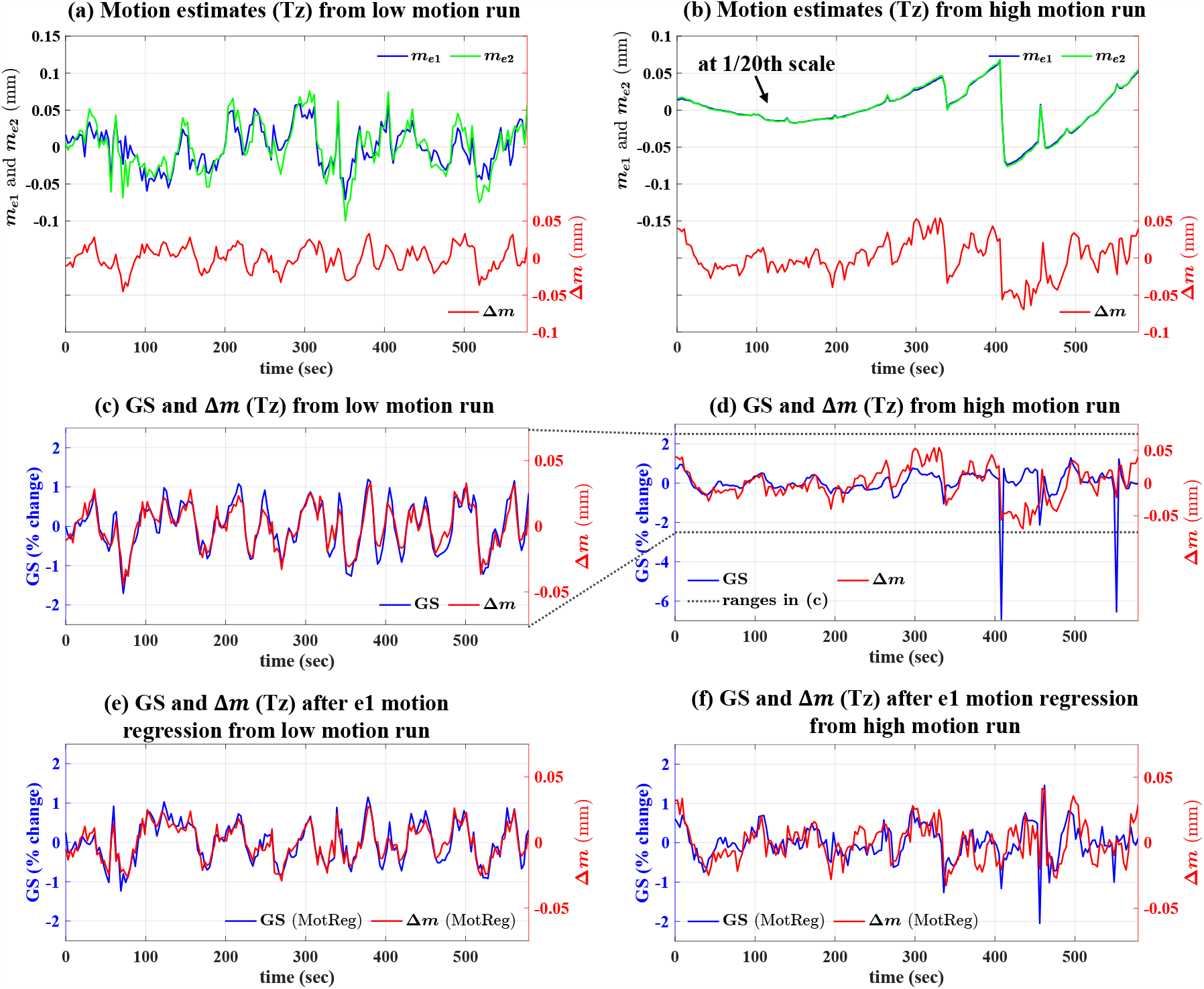
Motion estimates in the Tz axis, including ***m***_*e*1_ (blue), ***m***_*e*2_ (green) and Δ***m*** (red) from (a) low and (b) high motion runs. Panels (c) and (d) show the GS (blue) and Δ***m*** (red) from the low and high motion runs, respectively. Panels (e) and (f) show the GS (blue) and Δ***m*** (red) after motion regression (denoted as MotReg; with ***m***_*e*1_ as regressor) from the low and high motion runs, respectively.

Fig. 1 (c) and (d) show the GS and Δ***m*** from the low motion and high motion runs, respectively. For the low motion run, Δ***m*** covaries with the GS throughout the run, leading to a strong *r*(Δ***m***, GS) of 0.93. The high motion run shows a weaker *r*(Δ***m***, GS) of 0.34 as compared to the low motion run, and reflects motion artifacts in the GS. After motion regression (with ***m***_*e*1_; panels e and f), the *r*(Δ***m***, GS) for the high motion run increases to 0.62, while the *r*(Δ***m***, GS) of the low motion run remains at a high value of 0.92, suggesting that the relation between the GS and Δ***m*** is enhanced when motion artifacts are minimized.

### 3.2. Significance testing for r(Δ***m***, *GS*) *values*

To examine the presence of BOLD-weighted GS bias over runs and motion axes, we assessed the significance of *r*(Δ***m***, GS) values on a per-run and per-axis basis using permutation-based empirical null distributions. Fig. 2 shows two-sided violin plots of the distributions of *r*(Δ***m***, GS) values (blue) and the empirical null distributions (green) for all six motion axes (a) before and (b) after e1 motion regression. The blue solid lines and circles represent the median values for each distribution of measured *r*(Δ***m***, GS) values. The dashed lines represent the *r*(Δ***m***, GS) values corresponding to a Bonferroni corrected p-value threshold of 0.05 (two-sided) assessed from the empirical null distributions. The dark red square markers represent the percentages of runs showing *r*(Δ***m***, GS) values that are significantly different from zero.

**Figure 2:**
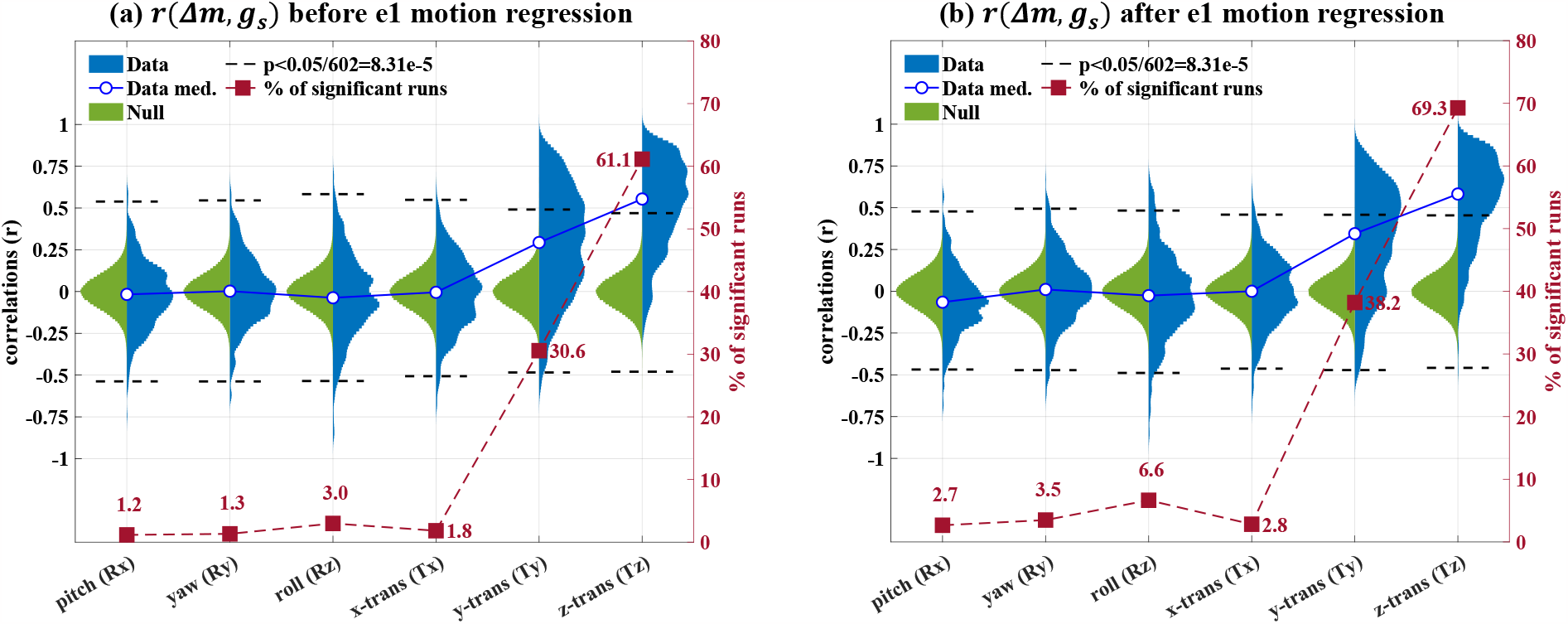
Two-sided violin plots showing the distributions of *r*(Δ***m***, GS) values (blue) and the empirical null distributions (green) for all six motion axes (a) before and (b) after e1 motion regression. The blue solid lines and circles represent the median values for each data distribution. For each motion axis, a Bonferroni-corrected run-wise p-value threshold of 0.05/602 was used, where 602 is the number of runs. The black dashed lines show *r*(Δ***m***, GS) values that correspond to the p-value thresholds (two-sided) assessed from the empirical null distributions. The dark red dashed lines and square markers represent the percent of the runs with significant *r*(Δ***m***, GS) values.

In the Tz axis, 61.1% and 69.3% of the runs show significant positive *r*(Δ***m***, GS) values before and after e1 motion regression, respectively. The group median *r*(Δ***m***, GS) value increases from 0.55 to 0.58 after e1 motion regression, with 96 out of 602 total runs showing *r*(Δ***m***, GS) values larger than 0.8 after e1 motion regression. In the Ty axis, 30.6% and 38.2% of the runs show significant *r*(Δ***m***, GS) values before and after e1 motion regression, respectively. The group median *r*(Δ***m***, GS) value increases from 0.29 to 0.34 after e1 motion regression. Fig. S1 and S2 show examples of the GS and the motion estimates, including ***m***_*e*1_, ***m***_*e*2_ and Δ***m*** in the Tz and Ty axes, respectively. Together, these findings indicate the presence of BOLD-weighted GS bias in the Tz and Ty motion estimates.

For the other motion axes (Rx, Ry, Rz and Tx), the percent of significant *r*(Δ***m***, GS) values fluctuates around 5%, ranging from 1.2% to 6.6%, indicating minimal BOLD-weighted GS bias for these axes.

### 3.3. Visualizing the spatial maps underlying r(Δ***m***, *GS*) *values*

In the previous section, BOLD-weighted GS bias in the motion estimates was identified by examining the temporal correlations between the GS and Δ***m***. Furthermore, as described in Eq. 4, *r*(Δ***m***, GS) can be approximated as the product of the difference of 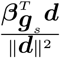 values from e2 and e1 and a scaling factor 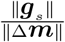. The approximation provides a unique angle to interpret the GS bias by looking at the relation of the 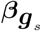 and ***d*** spatial maps. Fig. 3 visualizes 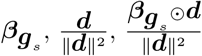, and the 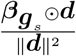 difference maps in the Tz and Tx axes from an example run, where ⊙ represents element-wise multiplication.

**Figure 3:**
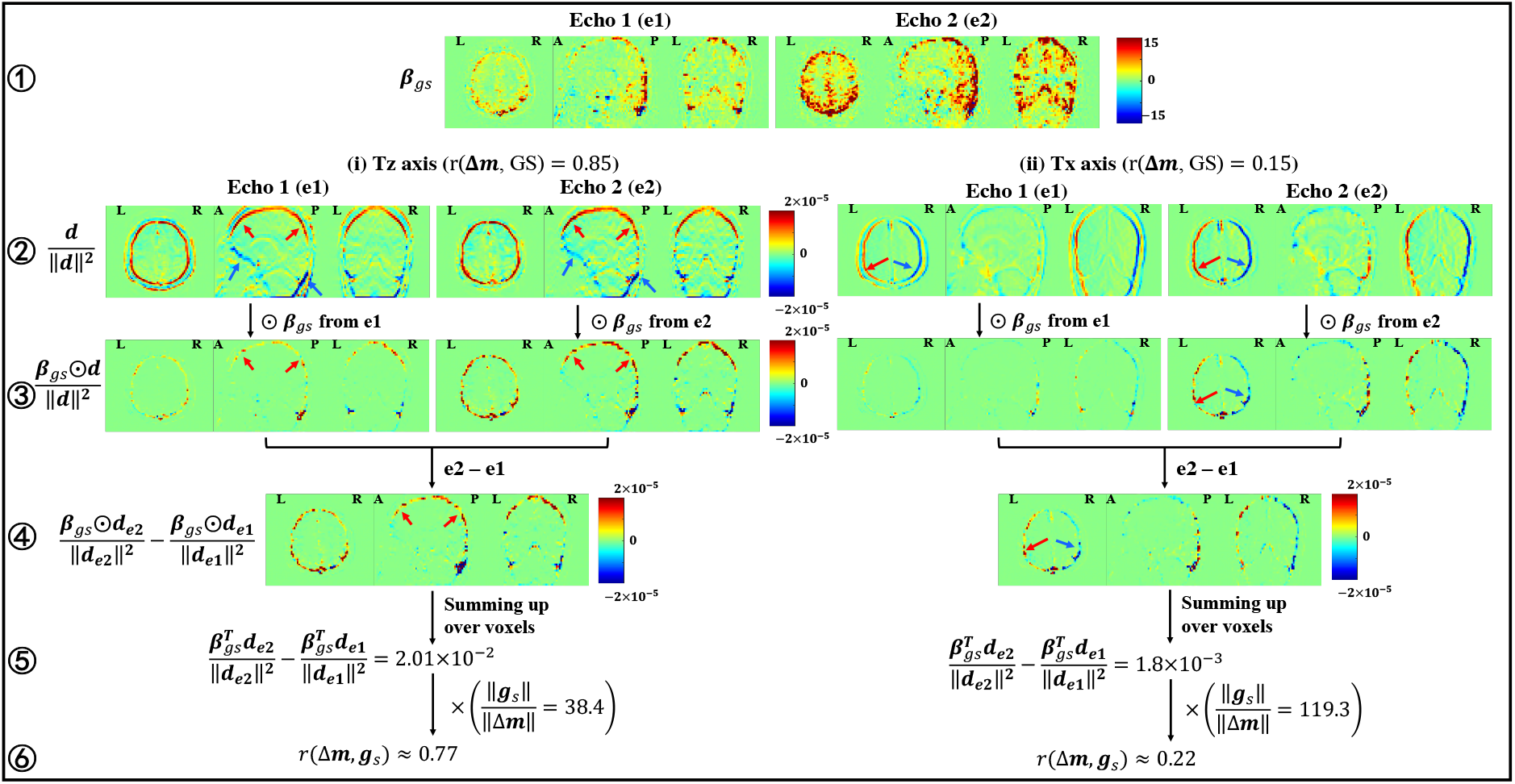
Visualization of the spatial maps underlying *r*(Δ***m***, GS) in the Tz (rows 2-6 on the left) and Tx (rows 2-6 on the right) axes from an example run. For each map, three representative slices (one axial, one sagittal and one coronal) are plotted. From top to bottom, rows 1 through 3 show the 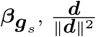, and 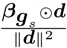 maps from e1 and e2, where ⊙ represents element-wise multiplication. Row 4 shows 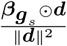 difference (e2-e1) maps. From left to right in rows 2 and 3, the first and second columns show the e1 and e2 maps in the Tz axis, and the third and fourth columns show the e1 and e2 maps in the Tx axis. The red and blue arrows point to brain regions showing high positive and negative values in the maps, respectively.

In the Tz axis, 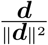 exhibits high positive and negative values at the superior and inferior edges of the brain, respectively, whereas 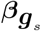 exhibits positive values across the cortex, including the superior edge of the brain, but exhibits relatively low values (approaching zero) along the inferior edge. As a result, most of the high values in 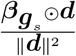 are positive with greater amplitudes for e2, resulting in high positive values in the 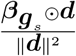 difference map. Consequently, summing up the 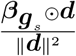 difference map leads to a high *r*(Δ***m***, GS) value. In contrast, for the Tx axis, 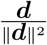 shows high positive and negative values on the left and right edges, respectively, whereas 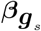 shows high positive values across the cortex, including both the left and right edges. The resulting high positive and negative values observed in the 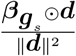 difference map tend to cancel out, resulting in a lower *r*(Δ***m***, GS) value. Together, these observations suggest that the GS bias in the Tz motion estimates results from 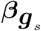 and ***d*** sharing a relatively similar top-bottom asymmetric spatial pattern.

In the Ty axis, as shown in Fig. S4, 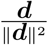 exhibits negative and positive values at the anterior and posterior edges of the brain, respectively, and 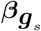 exhibits positive values across the cortex, including the posterior edge of the brain, but exhibits relatively low values (approaching zero) along the anterior edge. As a result, the positive values in the 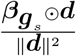 difference show relatively greater amplitudes as compared to the negative values, resulting in a high *r*(Δ***m***, GS) value.

Taken together, the GS-induced bias in the Ty and Tz motion estimates reflect the asymmetries in 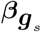 maps, which show higher values at the superior and posterior parts of the brain as compared to the inferior and anterior parts of the brain.

### 3.4. Effect of the bias on ROI-ROI FC via motion regression

In this section, we investigate how the presence of the GS-induced bias in the motion estimates affects the ROI-ROI FC by examining the differences in the ROI-ROI FC calculated after e2 and e1 motion regression. The e2-e1 differences in r-values and z-scores are denoted as Δ*r* and Δ*z*, respectively. Note that motion censoring was applied before motion regression. Fig. 4 shows Δ*r* and Δ*z* for all pairs of ROIs averaged over four groups of runs: (a) low motion and high aGS runs, (b) low motion and low aGS runs, (c) high motion and high aGS runs, and (d) high motion and low aGS runs. All four groups demonstrate negative Δ*r* and Δ*z* values for all pairs of ROIs, suggesting that the bias in the motion estimates can reduce FC estimates via motion regression. Comparing groups, we observed that the runs with low motion and high aGS exhibit the most negative Δ*r* and Δ*z* values (Δ*r* ranging from -0.02 to -0.09 and Δ*z* ranging from -0.63 to -1.41), while the runs with high motion and low aGS show Δ*r* and Δ*z* values close to zero (Δ*r* ranging from -0.00 to -0.02 and Δ*z* ranging from -0.05 to -0.33).

**Figure 4:**
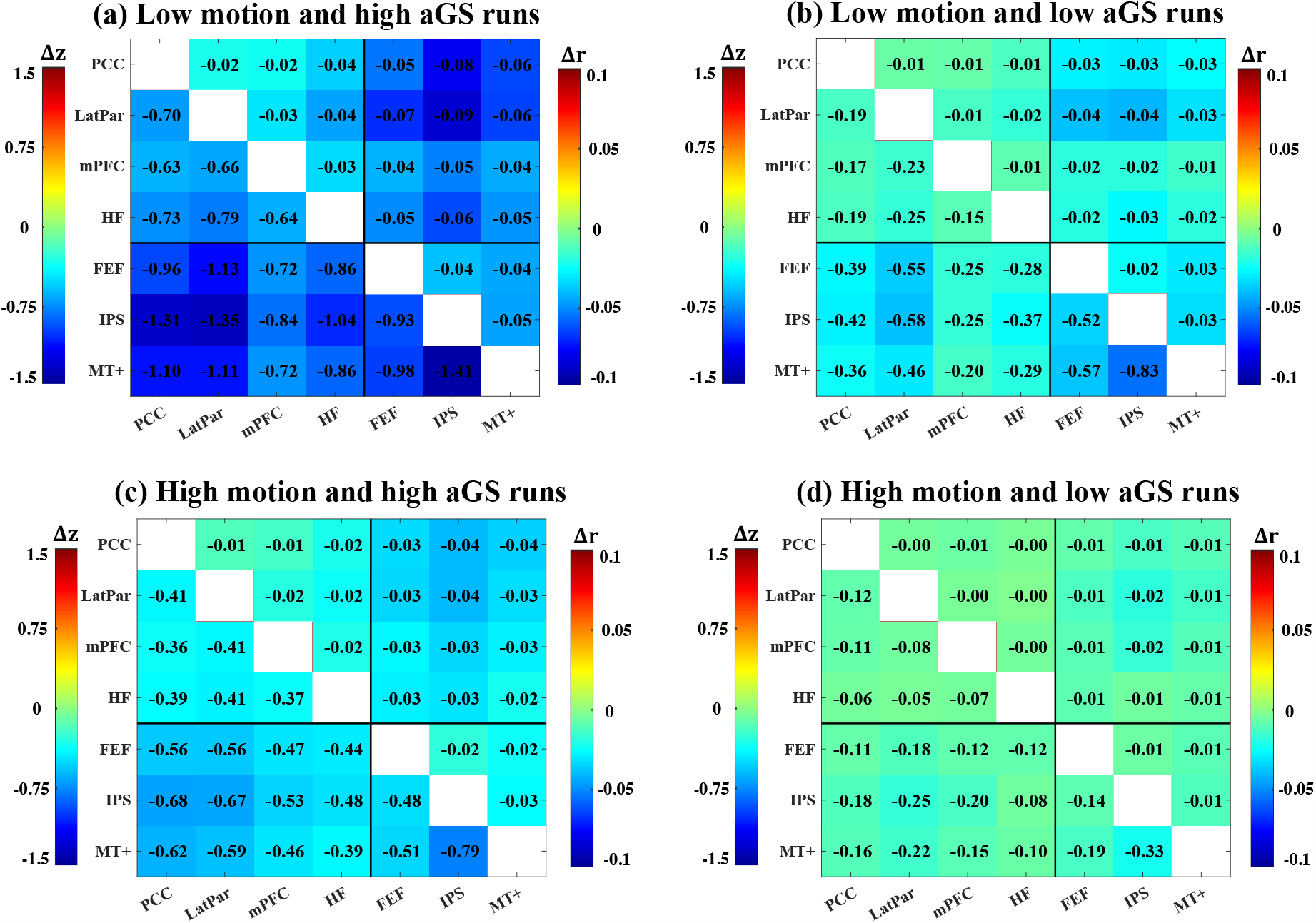
Differences (e2-e1) in ROI-ROI FC calculated after e2 and e1 motion regression. The differences were averaged across runs within each of four groups: (a) low motion and high aGS runs, (b) low motion and low aGS runs, (c) high motion and high aGS runs and (d) high motion and low aGS runs. Each subplot is divided into an upper right triangle showing the average differences in r-values and a lower left triangle showing the average differences in z-scores.

Furthermore, a one-way ANOVA on the mean Δ*z* values calculated over ROI pairs showed that there is a significant (*p* < 1 × 10^−6^, *F*_3,598_ = 40.53) difference among groups. The results from the post-hoc t-tests indicate that the low motion and high aGS runs exhibit significantly (*p* < 1 × 10^−4^) more negative mean Δ*z* values as compared to the other three groups, whereas the high motion and low aGS runs exhibit significantly (*p* < 1 × 10^−6^) less negative mean Δ*z* values as compared to the other three groups.

There is no significant (*p* = 0.0148 > 0.01) difference between the group of low motion and low aGS runs and the group of high motion and high aGS runs. The distributions of the mean Δ*z* values over ROI pairs for the four groups are shown in Fig. S6.

Fig. S7 and S8 demonstrate that the observed effect of the bias is dominated by the motion estimates in the Ty and Tz axes. In Fig. S7, when including only the Ty and Tz motion regressors, the results are similar to those shown in Fig. 4. In contrast, when excluding the Ty and Tz motion regressors (Fig. S8), we observed minimal differences between the FC estimates obtained with e1 and e2 motion regression. Together, these results indicate that regressing out the motion estimates with GS-induced bias may lead to reductions in FC estimates, with a stronger effect for runs with higher aGS and lower head motion.

#### 3.4.1. Effect of the bias on young vs. old group-level analysis

As shown in Fig. 5 (a) and (b), young subjects show significantly (*p* < 1 × 10^−6^) higher aGS and lower mean FD values as compared to the old subjects. Consequently, as shown in Fig. 5 (c), the young subjects show significantly (*p* < 1 × 10^−6^) more negative mean Δ*z* (e2-e1) values over ROI pairs as compared to the old subjects. The mean Δ*z* values over runs and ROI pairs are -0.69 and -0.17 for the young and old subjects, respectively. Furthermore, Fig. 5 (d) and (e) visualize the mean Δ*r* and Δ*z* values over the young and old subjects for all ROI pairs. We observed that for all ROI pairs, the young subjects show more negative Δ*r* and Δ*z* values as compared to the old subjects. Since Δ*z* and Δ*r* reflect the effect of the bias when performing motion regression, these results suggest a greater impact of the bias on the young subjects as compared to the old subjects.

**Figure 5:**
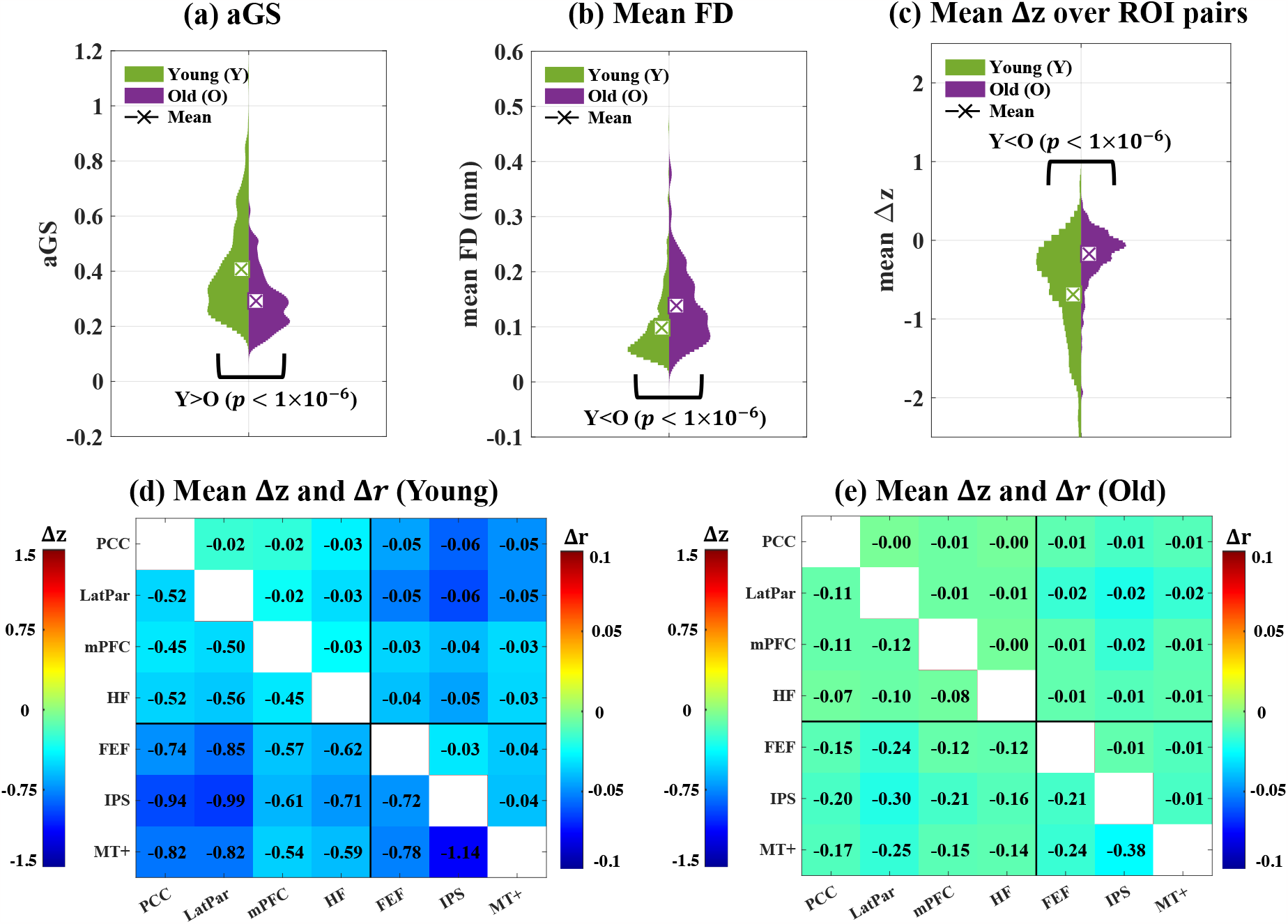
(a-c) Two-sided violin plots showing the distributions of (a) aGS, (b) mean FD and (c) mean Δ*z* (e2-e1) over ROI pairs for the young (green) and old (purple) subjects. Two-sided permutation tests were calculated to assess the significance thresholds for the differences between the young and old subjects. Young subjects show significantly (*p* < 1 × 10^−6^) larger aGS, smaller mean FD and more negative mean Δ*z* as compared to the old subjects. Panels (d-e) show the mean Δ*z* and Δ*r* over (d) young and (e) old subjects for all ROI pairs. Each subplot is divided into an upper right triangle showing the average differences in r-values and a lower left triangle showing the average differences in z-scores.

Moreover, we examined whether the bias affects the group-level analysis between the young and the old subjects. Fig. 6 (a) shows the differences in ROI-ROI FC between the young and old subjects calculated after motion censoring and e1 motion regression. We observed that the young subjects show significantly (*p* < 1 × 10^−3^) higher FC than the old subjects for all ROI pairs, except for the pair between IPS and LatPar. However, after e2 motion regression, as shown in Fig. 6 (b), the significant differences for five pairs of ROIs become insignificant (*p* > 0.01), including the pairs between (1) PCC and FEF, (2) PCC and IPS, (3) LatPar and FEF, (4) LatPar and MT+ and (5) FEF and IPS. Additionally, Fig. 6 (c) and (d) show that e2 motion regression reduces the FC differences between young and old for all pairs of ROIs as compared to e1 motion regression. Furthermore, as shown in Fig. S9 and S10, we verified that the observed effect is dominated by regressing out the Ty and Tz motion regressors. Together, these results demonstrate that motion regression with biased motion estimates can lead to underestimation of FC differences between the young and old subjects.

**Figure 6:**
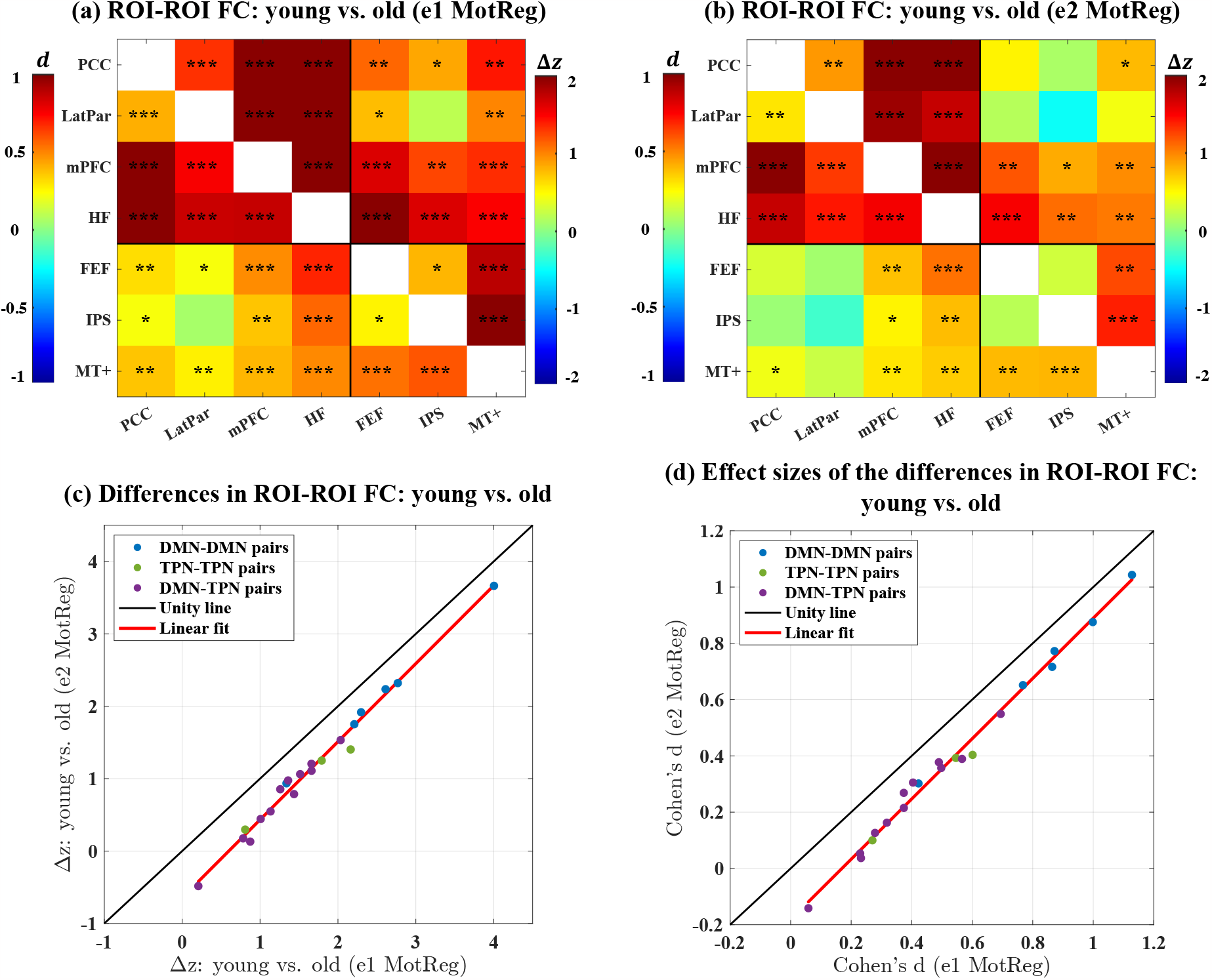
(a-b) ROI-ROI FC differences (young-old) between young and old subjects calculated after (a) e1 and (b) e2 motion regression. Each subplot is divided into an upper right triangle showing the averaged differences in z-scores between young and old subjects and a lower left triangle showing the effect size of the differences. The black asterisks indicate the statistical significance of the differences assessed by permutation tests (*: *p* < 0.01, **: *p* < 1 × 10^−3^, ***: *p* < 1 × 10^−6^). A positive value (red color) indicates that the young subjects show higher connectivity as compared to the old subjects. (c) Scatter plot comparing the differences in ROI-ROI FC between the young and the old subjects calculated after e1 and e2 motion regression (denoted as e1 MotReg and e2 MotReg, respectively) for all pairs of ROIs. (d) Scatter plot comparing the effect sizes of the differences in ROI-ROI FC between the young and the old subjects calculated after e1 and e2 motion regression for all pairs of ROIs.

## 4. Discussion

In this study, we used a public resting-state MEfMRI dataset to examine whether global brain activity can lead to bias in rsfMRI motion estimates and to characterize the potential impact on rsFC estimates. By examining the correlation between the GS and the difference in the motion estimates from the first and second echoes, we found evidence for GS-related bias in the Tz and Ty motion estimates. We also demonstrated that the GS-induced bias can lead to underestimation of rsFC estimates when using motion regression, with low motion and high aGS runs exhibiting the greatest reductions in FC due to the bias and high motion and low aGS runs showing minimal reductions in FC. Finally, we showed that regression with biased motion estimates can reduce rsFC differences between groups of young and old subjects, due in part to different levels of aGS and head motion between the groups.

Extending prior studies that examined bias in task-based fMRI motion estimates, we identified BOLD-weighted bias in rsfMRI motion estimates and investigated its effect on rsFC. Moreover, utilizing multi-echo fMRI data, we proposed a novel method to detect the BOLD-weighted bias in real motion estimates over a large sample of runs with rigorous statistical tests. Furthermore, to investigate the underlying cause of the observed bias, we proposed an empirical approximation to *r*(Δ***m***, GS) and used this approximation to demonstrate how the presence of the GS-induced bias in the motion estimates may be attributed to the superior-inferior and posterior-anterior asymmetric spatial patterns in the GS beta coefficient map 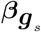. In prior work (Freire and Mangin, 2001; Freire et al., 2002), the existence of the bias in task-based fMRI was primarily illustrated using simulations and the use of experimental data was limited to one scan.

While the interpretation of the GS is still controversial (reviewed in (Liu et al., 2017)), there is growing evidence suggesting that the GS is linked to vigilance (also known as arousal level) (Wong et al., 2013, 2016; Falahpour et al., 2016, 2018; Liu et al., 2018). For example, Falahpour et al. (2016) found that the GS is negatively correlated with EEG measures of vigilance. Furthermore, studies have shown that a vigilance template, calculated from voxel-wise correlations between the EEG vigilance measures and the fMRI signal, can be used to estimate vigilance fluctuations in fMRI scans (Chang et al., 2016; Falahpour et al., 2018; Goodale et al., 2021). As shown in Supplementary Material, the superior-inferior and posterior-anterior asymmetric spatial patterns in the 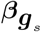 maps show a significant spatial correlation with the patterns observed in the vigilance template, suggesting that the asymmetric patterns may partly reflect spatial variations in the effect of vigilance variations on the fMRI signal. Furthermore, as described in Supplementary Material, image intensity variations due to factors such as inhomogeneities in the receive coil sensitivity patterns (Belaroussi et al., 2006) are another potential source of the posterior-anterior asymmetric pattern in 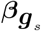.

In this work, we performed a detailed analysis of the bias in motion estimates obtained with AFNI *3dvolreg*, which is widely used for preprocessing of fMRI data. As described in Supplementary Material, a preliminary examination shows that bias is also observed when using other software packages, such as SPM (Friston et al., 1995), FSL (Jenkinson and Smith, 2001; Jenkinson et al., 2002), and ANTS (Avants et al., 2011, 2014). The bias observed with SPM is similar to that observed with AFNI, while FSL and ANTS show relatively lower levels of bias. Future work is needed to thoroughly examine how the choice of algorithm (and associated cost functions) affects the level of the bias in the motion estimates. In addition, future work focused on developing registration algorithms with reduced bias would be of interest.

In this study, we demonstrated that regression with biased motion estimates can reduce differences in rsFC between the young and old subjects, as assessed with ROIs in the DMN and TPN. Importantly, we showed that the significant rsFC differences for five pairs of ROIs became insignificant (*p* > 0.01) due to regression with the biased motion estimates, with four pairs corresponding to connections between the DMN and TPN. These findings suggest that regression with biased motion estimates may impede the detection of rsFC differences between the young and old groups. Because rsFC of the DMN plays a key role in understanding the aging brain (reviewed in (Ferreira and Busatto, 2013)), it can be important for future aging studies to minimize the effect of the bias on rsFC analyses. Based on our results, investigators can use motion estimates from the first echo data (if available) or consider excluding the Ty and Tz motion estimates from the motion regression. Moreover, our findings indicate that the biased motion estimates may lead to discrepancies in rsFC results via regression. Therefore, when comparing results from existing aging studies with different processing methodologies, the potential effect of the bias needs to be considered. Concerning this matter, future work that further examines the effect of the bias on the young vs. old differences in rsFC measures would be of interest.

Our findings revealed that the effect of the bias on the young vs. old rsFC analysis may be caused by different levels of aGS and head motion between the groups, with the old subjects showing a lower level of aGS and a higher level of head motion than the young subjects. The higher levels of motion in the old group have been consistently reported in prior rsfMRI studies (Saccà et al., 2021; Hausman et al., 2022), which reflects declines in executive functioning with aging (Hausman et al., 2022). A direct comparison of aGS between young and old groups does not appear to have been addressed in prior studies. However, previous studies have reported that old subjects show lower rsFC and BOLD variability (measured as the standard deviation of BOLD timeseries) in large-scale brain networks, including the DMN, compared to young subjects (Ferreira and Busatto, 2013; Grady and Garrett, 2018; Nomi et al., 2017; Kumral et al., 2020), supporting the observed lower aGS in the old subjects.

## 5. Conclusion

In conclusion, we found that resting-state global brain activity can lead to bias in Ty and Tz motion estimates obtained with a widely used motion correction algorithm. Furthermore, the bias in the motion estimates can lead to reductions in the rsFC measures obtained after motion regression, with an increasing level of bias for runs showing higher global signal amplitude and smaller head motion levels. This GS-related decrease in rsFC values is similar to the reduction seen with global signal regression (Liu et al., 2017), and therefore concerns about the effects of global signal regression may also apply when regressing out motion parameters estimated from rsfMRI data acquired at typical echo times (e.g. 30 ms). Moreover, our results show that regression with biased motion estimates can reduce group-level rsFC differences between young and old subjects. Similar effects may also be present in other rsfMRI studies in which the groups exhibit different mean levels of global signal amplitude. Our results suggest that a greater degree of caution should be used when interpreting rsFC differences obtained when motion regression is used in the processing of rsfMRI data.

## 6. Ethics Statement

The study that collected the open-source dataset was approved by the Institutional Review Board at Cornell University and the Research Ethics Board at York University. Written informed consent was obtained from all study participants.

## 7. Data and Code Availability

The dataset can be downloaded from OpenNeuro (dataset ds003592). Analysis code and files to generate the figures and results presented in this paper will be made available upon publication through the Open Science Framework DOI 10.17605/OSF.IO/SH79V.

## 8. Author Contributions

Y.M. and T.L. designed the experiments. Y.M. carried out the experiments and prepared the results. Y.M. and T.L. wrote and edited the main manuscript text. All authors discussed the results and provided interpretations of the results. All authors reviewed and provided feedback on the manuscript. All authors contributed to the article and approved the submitted version.

## Appendix A. Motion estimation in AFNI *3dvolreg*

In this section, we review the theory and implementation of AFNI *3dvolreg*. Let ***Y*** = [***y***_1_, ***y***_2_, …, ***y***_*K*_] ∈ ℝ^*N* ×*K*^ be the functional data, where *N* is the number of voxels, *K* is the number of volumes and ***y***_*k*_ ∈ ℝ^*N* ×1^ is the *k*th column of ***Y***, i.e. the *k*th volume of the data with *k* = 1, 2, …, *K*. Let ***y***_*r*_ ∈ ℝ^*N* ×1^ be the reference image used in the registration. To align ***y***_*k*_ to ***y***_*r*_, the algorithm first estimates the motion parameters ***a***_*k*_ ∈ ℝ^6×1^ (consisting of 3 rotational and 3 translational motion parameters) by iteratively minimizing the weighted least square cost function:

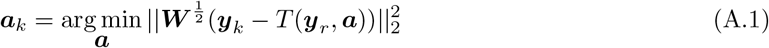

where ***W*** is a diagonal weight matrix and *T* : ℝ^*N* ×1^ → ℝ^*N* ×1^ is the function that performs rigid body transformation. Because we disabled the weights, we have ***W*** = ***I***, where ***I*** is the identity matrix. After obtaining ***a***_*k*_, the algorithm then transforms ***y***_*k*_ with −***a***_*k*_, which can be written as *T* (***y***_*k*_, −***a***_*k*_), to register ***y***_*k*_ to ***y***_*r*_. To solve the minimization problem in Eq. A.1, the algorithm linearizes the rigid body transformation with a first-order Taylor approximation, which can be written as

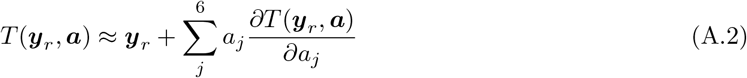

where *a*_*j*_ is the *j*th element of ***a***. Furthermore, the partial derivatives in Eq. A.2 are approximated with finite differences:

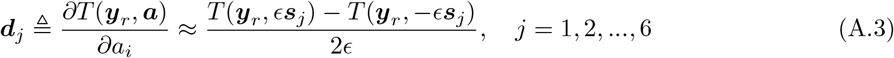

where *ϵ* is a scalar with small magnitude and ***s***_*j*_ is a 6 × 1 vector whose *j*th element is one while the other elements are zeros. Here, we use ***d***_*j*_ ∈ ℝ^*N* ×1^ to represent the partial derivative with respect to (w.r.t.) *a*_*j*_ and refer to it as the spatial derivative image w.r.t. the *j*th motion axis. The linearization of the transformation can then be written as

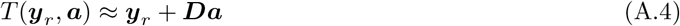

where ***D*** = [***d***_1_, ***d***_2_, …, ***d***_6_] ∈ ℝ^*N* ×6^. With the above linearization and ***W*** = ***I***, the minimization problem in Eq. A.1 can be written as

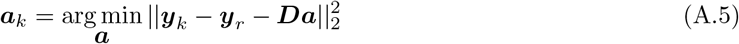

In AFNI *3dvolreg*, the motion parameters are estimated by iteratively minimizing the cost function in Eq. A.5. Let ***a***_*k,n*_ denote the motion parameters estimated at the *n*th iteration,

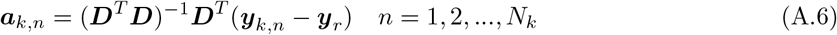

where *N*_*k*_ is the total number of iterations for the *k*th volume and ***y***_*k,n*_ can be written as

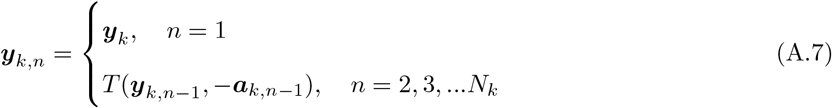

The algorithm stops the iterative process when ***a***_*k,n*_ is smaller than a fixed threshold (for the dataset used in this study, the translational and rotational thresholds are 0.03 mm and 0.02 degree, respectively) or *N*_*k*_ exceeds the maximum number of iterations, denoted as *N*_*max*_. The algorithm uses this iterative process to deal with the approximation error in the linearization of the rigid transformation described in Eq. A.4. In practice, for most volumes, the iterative process stops at the second iteration, suggesting that the parameters estimated in the second iteration are smaller than the thresholds and Eq. A.4 is a valid approximation. After the iterative estimation process stops, the motion parameters of the *k*th volume are calculated by summing the estimated parameters over all iterations:

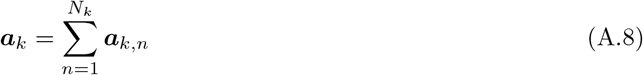

Stacking the motion parameters over volumes yields

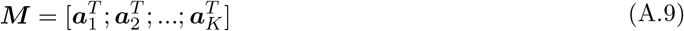

Here, the *j*th column of ***M*** represents the time series of the motion parameters of the *j*th motion axis (denoted as ***m***_*j*_). Furthermore, in order to explicitly show the relation between ***M*** and the functional data ***Y***, we define ***y***_*k,n*_ = ***y***_*r*_ for *N*_*k*_ < *n* ≤ *N*_*max*_ so that ***a***_*k,n*_ = **0** for *N*_*k*_ < *n* ≤ *N*_*max*_. Therefore, the motion parameters can be written as

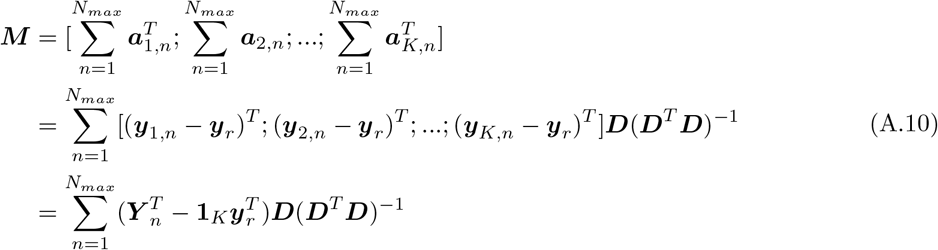

where ***Y*** _*n*_ = [***y***_1,*n*_, ***y***_2,*n*_, …, ***y***_*K,n*_] and **1**_*K*_ is a *K* by 1 all-one vector. Here, 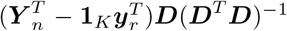 represents the parameters estimated at the *n*th iteration.

In practice, the algorithm directly estimates and outputs the transformation parameters used to align ***y***_*k*_ to ***y***_*r*_ (i.e., −***a***_*k*_) by using −***D***. Therefore, in this work, the transformation parameters generated by AFNI *3dvolreg* were multiplied by −1 to represent the motion parameters ***a***_*k*_.

## Appendix B. Empirical approximations to the motion estimates

To explore the key terms underlying *r*(Δ***m***, GS), we derived the following empirical approximations of the motion estimates:

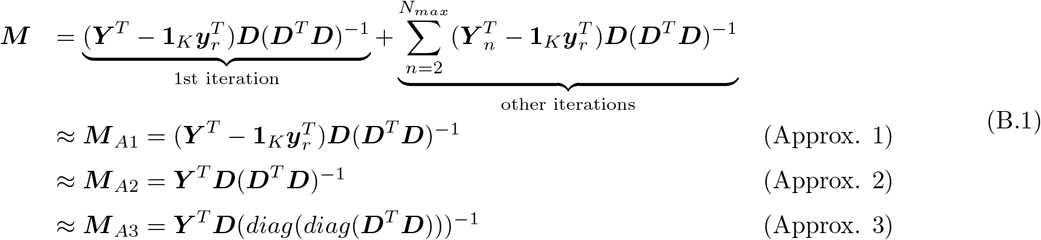

where **1**_*K*_ ∈ ℝ^*K*×1^ is an all-one vector and *diag()* denotes the Matlab function `diag()`. The first approximation of the motion estimates, denoted as ***M*** _*A*1_, represents the motion parameters estimated at the first iteration. This approximation is valid when Eq. A.4 Then, because the reference image is almost orthogonal to the derivative images (i.e., 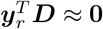), we can make the second approximation, denoted as ***M*** _*A*2_, which represents the motion parameters estimated without subtracting the reference image from the functional data. The third approximation of the motion estimates, denoted as ***M*** _*A*3_, represents the motion parameters estimated after zeroing out the off-diagonal terms in ***D***^*T*^ ***D***, reflecting the fact that the covariance between the derivative images in different axes is relatively small compared to the variance of the derivative images. To evaluate the approximations, the temporal correlations between ***M*** and the approximated motion estimates were calculated for each motion axis. As shown in Fig. S14, ***M*** is highly correlated with ***M*** _*A*1_ (mean *r* = 0.99), ***M*** _*A*2_ (mean *r* = 0.99), and ***M*** _*A*3_ (mean *r* = 0.91)). Fig. S15 displays the motion estimates before and after the approximations in the Tz and Ty axes from three example runs.

With the third approximation, the motion estimates of one motion axis, denoted as ***m***_*A*3_ ∈ ℝ^*K*×1^ can be written as

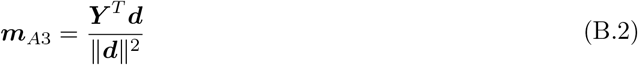

where ***d*** ∈ ℝ^*N* ×1^ is the spatial derivative image w.r.t. that motion axis.

## Appendix C. The approximation to *r*(Δ*m*, GS)

To interpret *r*(Δ***m***, GS), we derived a mathematical approximation that reveals the key factors. We begin with the expression

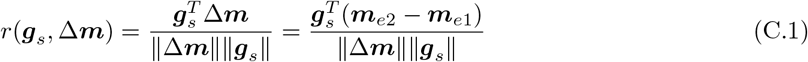

where ***g***_*s*_ ∈ ℝ^*K*×1^, Δ***m*** = ***m***_*e*2_ − ***m***_*e*1_ ∈ ℝ^*K*×1^ represent the GS and the difference between the e2 and e1 motion estimates, respectively, and ***m***_*ei*_ ∈ ℝ^*K*×1^ represents the motion estimates from the *i*th echo. Then, using the approximation described in Eq. B.2, we can write

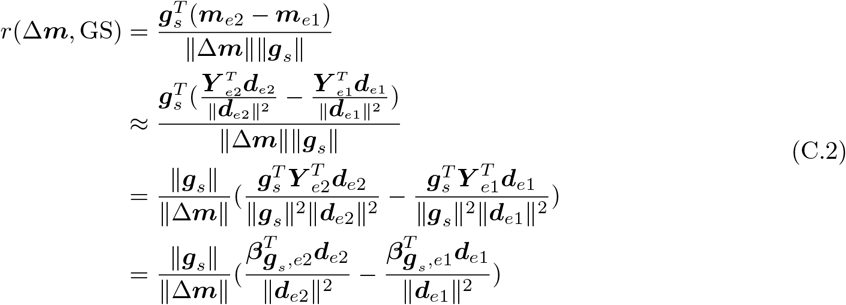

where 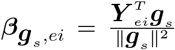 is the GS beta coefficient map calculated from the linear fit of the GS to the unregistered functional data of the *i*th echo.

