## Supplementary Materials for "Resting state global brain activity induces bias in fMRI motion estimates"

### Supplementary Material for "Resting state global brain activity induces bias in fMRI motion estimates"

#### S1. Supplementary Methods

##### *S1.1. Spatial correlation between the vigilance template and* $\boldsymbol{\beta}_{\boldsymbol{g}_s}$

To investigate whether the asymmetric spatial patterns in $\boldsymbol{\beta}_{\boldsymbol{g}_s}$ are associated with vigilance-related rsfMRI signal fluctuations, we compared $\boldsymbol{\beta}_{\boldsymbol{g}_s}$ from the example run shown in Fig. 3 to the eyes-open vigilance template estimated in (Falahpour et al., 2018). Here, $\boldsymbol{\beta}_{\boldsymbol{g}_s}$ was calculated on the functional data after registration, transformation to MNI152 standard space, and spatial smoothing with a 4mm FWHM Gaussian kernel. The vigilance template in (Falahpour et al., 2018) was in the N27 template, so it was transferred to the MNI152 standard space in this work. The spatial correlation between the vigilance template and the example $\boldsymbol{\beta}_{\boldsymbol{g}_s}$ was calculated.

##### *S1.2. Examining the intensity inhomogeneity in the fMRI data*

Prior work has shown that the spatial intensity inhomogeneity (also known as the bias fields) in fMRI data can lead to registration errors (Gonzalez-Castillo et al., 2013). We found that the images in the dataset exhibited posterior-anterior intensity inhomogeneity, which may contribute to the posterior-anterior asymmetry pattern observed in $\boldsymbol{\beta}_{\boldsymbol{g}_s}$. To illustrate the effect of the intensity inhomogeneity, we picked two low motion and high aGS runs as examples, one with a high Ty $r(\Delta\boldsymbol{m}, \text{GS})$ value of 0.87 (the example run shown in Fig. S4) and another one with a low Ty $r(\Delta\boldsymbol{m}, \text{GS})$ value of 0.07 (this example run was not shown before). For each run, the spatial intensity inhomogeneity was estimated using the reference image (i.e., the first volume) with the segmentation function provided in SPM (Gispert et al., 2004).

##### *S1.3. Comparing the GS-induced bias in motion estimates from AFNI, SPM, FSL and ANTS*

In addition to AFNI *3dvolreg*, we also performed a preliminary examination of the $r(\Delta\boldsymbol{m}, \text{GS})$ values calculated with the motion estimates from three other commonly used algorithms: SPM *spm_realign* (Friston et al., 1995), FSL *mcflirt* (Jenkinson and Smith, 2001; Jenkinson et al., 2002), and ANTS *antsMotionCorr* (Avants et al., 2011, 2014).

For SPM *spm_realign* and FSL *mcflirt*, we generally used the default settings while making a few necessary modifications. For SPM, we disabled the default spatial smoothing of 5 mm FWHM by setting the parameter *fwhm* to 0, since we found that applying spatial smoothing before registration can increase the level of the GS-induced bias in the motion estimates. Also, we disabled registration to the mean volume by setting the parameter *rtm* to 0. For FSL, we changed the reference volume to the first volume to be consistent with AFNI *3dvolreg*. As FSL supports different cost functions, in addition to the default cost function (normalized correlation, NC), we also estimated the motion parameters using the least-squares (LS) and mutual information (MI) cost functions. The LS cost function is used by AFNI and SPM, and the MI cost function has demonstrated the ability to reduce the neural-related bias in motion estimates (Freire and Mangin, 2001; Freire et al., 2002).

For ANTS *antsMotionCorr*, there is no default setting. Therefore, we followed the settings in an example that uses a rigid transformation and the MI cost function (github link). The first image was used as the reference image. It is worth mentioning that the sampling percentage was set to 10%, meaning that the algorithm only samples and uses 10% of the voxels during motion estimation. To make a fair comparison with the other algorithms, we also examined the motion parameters estimated using all the voxels (i.e., setting the sampling percentage to 100%).

For each run, the motion was estimated using both the e1 and e2 data. Because different algorithms have different default orientations, they output motion parameters in different orientations. To account for these differences, we multiplied the motion parameters by either 1 or -1 so that the resulting motion parameters were in the RAI orientation. The mean and the linear and quadratic trends were regressed out from the motion estimates.

Following the descriptions in Methods 2.4, for each run and motion axis, $r(\Delta\boldsymbol{m}, \mathrm{GS})$ was calculated and the significance of the $r(\Delta\boldsymbol{m}, \mathrm{GS})$ values was assessed by empirical null distributions. The mean $r(\Delta\boldsymbol{m}, \mathrm{GS})$ values over runs and the percent of runs showing significant $r(\Delta\boldsymbol{m}, \mathrm{GS})$ values were calculated to assess the level of the GS-induced bias in the motion estimates.

#### S2. Supplementary Results

##### *S2.1. $r(\Delta\boldsymbol{m}, GS)$ calculated using motion estimates with AFNI default weightings and GS after registration*

In the main text, when we calculated $r(\Delta\boldsymbol{m}, \mathrm{GS})$, we used the motion parameters estimated without AFNI default weightings and the GS calculated before volume registration. Here, we examined how these choices affect the $r(\Delta\boldsymbol{m}, \mathrm{GS})$ values. Fig. S3 shows comparisons of (a) mean $r(\Delta\boldsymbol{m}, \mathrm{GS})$ over runs and (b) the percentage of runs showing significant $r(\Delta\boldsymbol{m}, \mathrm{GS})$ values calculated with the motion parameters estimated with and without weightings (denoted as $\mathrm{Mot}_{\mathrm{wt}}$ and Mot, respectively) and the GS calculated before and after volume registration (denoted as GS and $GS_{vr}$, respectively). Results obtained with GS and $GS_{vr}$ exhibit similar values for both the mean $r(\Delta\boldsymbol{m}, \mathrm{GS})$ and the percentage of runs showing significant $r(\Delta\boldsymbol{m}, \mathrm{GS})$. Comparing motion estimates with and without default weightings, we observe that $\mathrm{Mot}_{\mathrm{wt}}$ shows higher percentages of runs showing significant $r(\Delta\boldsymbol{m}, \mathrm{GS})$ values in all motion axes except for the Ty axis. In the Ty axis, $\mathrm{Mot}_{\mathrm{wt}}$ shows a slight decrease (about 1%) in the percentage of runs showing significant $r(\Delta\boldsymbol{m}, \mathrm{GS})$ values. These results suggest that using AFNI default weightings may lead to a slightly higher level of the bias in the motion estimates in all motion axes except for the Ty axis.

##### *S2.2. Effect of the bias on ROI-ROI FC via motion regression without motion censoring*

The differences in the ROI-ROI FC calculated after e2 and e1 motion regression (i.e., $\Delta z$) without motion censoring are plotted in Fig. S5 and compared to $\Delta z$ calculated with motion censoring in Fig. S6. In Fig. S5, we observe that the range and pattern of the average $\Delta z$ values obtained without censoring are similar to those of the average $\Delta z$ values obtained with censoring shown in Fig. 4.

Fig. S6 shows two-sided violin plots of the distributions of mean $\Delta z$ (e2-e1) over ROI pairs calculated with (blue) and without (green) motion censoring for the four groups of runs. For each group, a paired t-test was calculated to compare the mean $\Delta z$ with and without motion censoring. The mean $\Delta z$ without motion censoring is significantly ($p < 1 \times 10^{-3}$) less negative than the mean $\Delta z$ with motion censoring for the low motion and high aGS group and the low motion and low aGS group. Although the differences in the mean $\Delta z$ values are significant, the magnitudes of the differences are relatively small as compared to the magnitudes of the mean $\Delta z$ values. For the low motion and high aGS group, mean $\Delta z = -0.93$ and $-0.87$ with and without motion censoring, respectively, resulting in a difference of $-0.06$. The effect size of the difference is -0.29. For the low motion and low aGS group, mean $\Delta z = -0.36$ and $-0.33$ with and without motion censoring, respectively, resulting in a difference of $-0.03$. The effect size of the difference is -0.30. No significant ($p > 0.01$) differences were found in the other groups.

Taken together, these results suggest that motion censoring has little impact on the changes in rsFC estimates due to regression with biased motion estimates.

##### *S2.3. Spatial correlation between the vigilance template and $\boldsymbol{\beta}_{\boldsymbol{g}_s}$*

Fig. S11 compares $\boldsymbol{\beta}_{\boldsymbol{g}_s}$ from the example run shown in Fig. 3 to the eyes-open vigilance template estimated in (Falahpour et al., 2018) (both spatial maps were transferred to the MNI152 standard space).

As shown in Fig. S11 (b), the estimated vigilance template exhibits similar superior-inferior and posterior-anterior asymmetric spatial patterns to the example $\boldsymbol{\beta}_{\boldsymbol{g}_s}$ map, resulting in a strong negative correlation ($r = -0.38$, $p < 1 \times 10^{-6}$) between $\boldsymbol{\beta}_{\boldsymbol{g}_s}$ and the vigilance template. The observed negative correlation is consistent with the negative correlations between the GS and the EEG vigilance found in (Falahpour et al., 2016, 2018). Hence, the asymmetric spatial patterns in $\boldsymbol{\beta}_{\boldsymbol{g}_s}$ may be linked to resting-state global brain activity associated with vigilance changes.

##### *S2.4. Examining the intensity inhomogeneity in the fMRI data*

Fig. S12 shows the reference images and estimated bias field maps from the example runs. For the example run with a strong $r(\Delta\boldsymbol{m}, \mathrm{GS})$ in the Ty axis shown in Fig. S12 (a), its reference image and estimated bias field map exhibit higher values at the posterior part of the brain and lower values at the anterior part of the brain. In contrast, for the run with a Ty $r(\Delta\boldsymbol{m}, \mathrm{GS})$ value close to zero (shown in Fig. S12 (b)), its reference image and estimated bias field map show comparable values at the anterior and posterior parts of the brain. These results suggest that the intensity inhomogeneity in the fMRI data used for this study may contribute to the posterior-anterior asymmetric pattern in $\boldsymbol{\beta}_{\boldsymbol{g}_s}$. In addition, although both runs showed high positive $r(\Delta\boldsymbol{m}, \mathrm{GS})$ values in the Tz axis, their estimated bias field maps showed minimal superior-inferior asymmetry, suggesting that the intensity inhomogeneity is not contributing to the superior-inferior asymmetry in $\boldsymbol{\beta}_{\boldsymbol{g}_s}$. Because spatial intensity inhomogeneity in fMRI data is affected by multiple factors, including radio-frequency coil homogeneity, the pulse sequences used, and the imaged object itself (Belaroussi et al., 2006), the impact of the intensity inhomogeneity on motion bias may vary greatly across datasets.

##### *S2.5. Assess the GS-induced bias in motion estimates from SPM, FSL and ANTS*

Fig. S13 (a) and (b) show the mean $r(\Delta\boldsymbol{m}, \mathrm{GS})$ values and the percent of runs showing significant $r(\Delta\boldsymbol{m}, \mathrm{GS})$ values, respectively. For all algorithms and cost functions, the Ty and Tz axes exhibit higher mean $r(\Delta\boldsymbol{m}, \mathrm{GS})$ values and a higher percentage of runs showing significant $r(\Delta\boldsymbol{m}, \mathrm{GS})$ values as compared to the other motion axes, consistent with the results in the main text. As the mean $r(\Delta\boldsymbol{m}, \mathrm{GS})$ values and the percent of runs showing significant $r(\Delta\boldsymbol{m}, \mathrm{GS})$ values follow a similar pattern, we will focus on the mean $r(\Delta\boldsymbol{m}, \mathrm{GS})$ values.

In the Tz axis, when comparing different cost functions, the LS cost function shows higher mean $r(\Delta\boldsymbol{m}, \mathrm{GS})$ values as compared to the NC and MI cost functions. When comparing across algorithms,

AFNI (LS) and SPM (LS) show comparably high mean $r(\Delta\boldsymbol{m}, \mathrm{GS})$ values, FSL with different cost functions (LS, NC and MI) show moderate mean $r(\Delta\boldsymbol{m}, \mathrm{GS})$ values, and ANTS (MI) shows the lowest mean $r(\Delta\boldsymbol{m}, \mathrm{GS})$ values. It is worth noting that for ANTS (MI), using 10% of the voxels during registration shows a lower mean $r(\Delta\boldsymbol{m}, \mathrm{GS})$ value as compared to using all the voxels. In the Ty axis, AFNI (LS) shows the highest mean $r(\Delta\boldsymbol{m}, \mathrm{GS})$ value, SPM (LS) and FSL with different cost functions show moderate mean $r(\Delta\boldsymbol{m}, \mathrm{GS})$ values, and ANTS (MI) shows the lowest mean $r(\Delta\boldsymbol{m}, \mathrm{GS})$ values. Overall, we observe that both the software package and cost function affect the level of the bias, with differences in the software implementation of the same cost function playing a significant role.

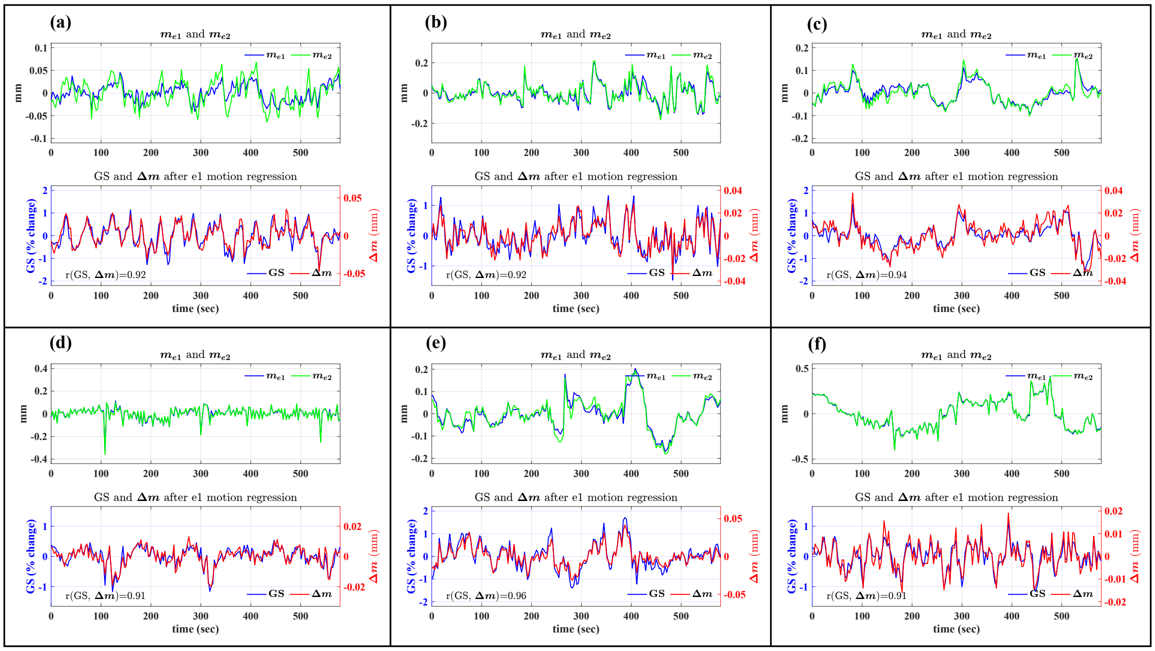


Figure S1: Global signal (GS) and motion estimates in the Tz axis, including $\boldsymbol{m}_{e1}$, $\boldsymbol{m}_{e2}$ and $\Delta \boldsymbol{m}$ from six example runs showing high $r(\Delta \boldsymbol{m}, \mathrm{GS})$. The e1 motion regressors were regressed out from the GS and $\Delta \boldsymbol{m}$.

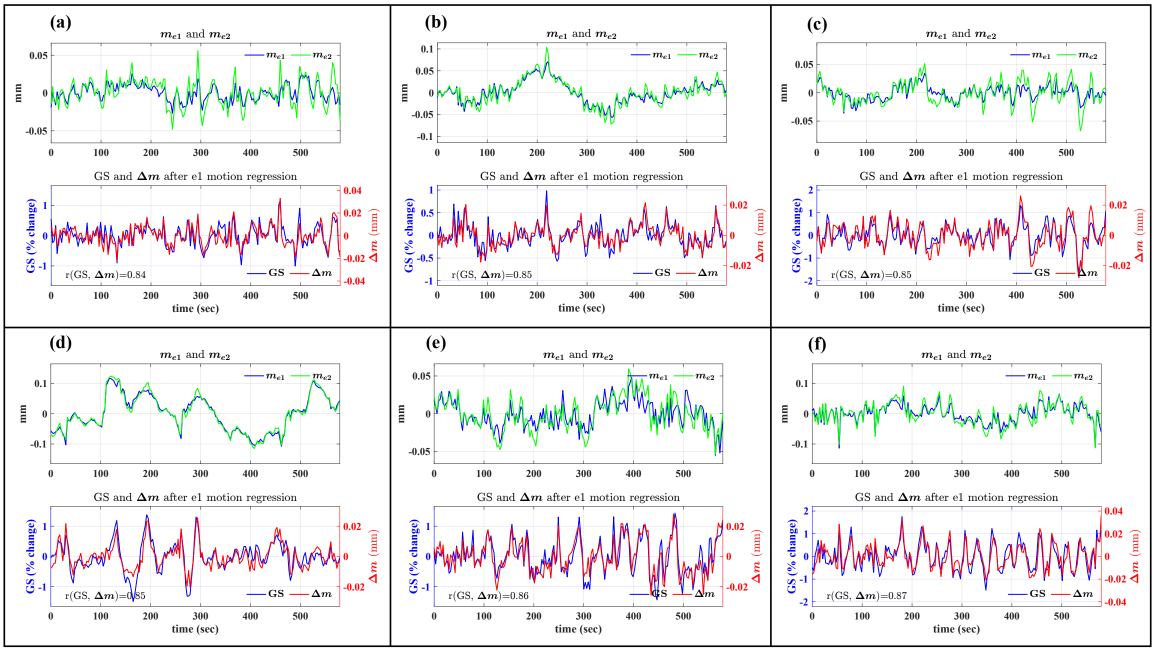

Figure S2: Global signal (GS) and motion estimates in the Ty axis, including $\boldsymbol{m}_{e1}$, $\boldsymbol{m}_{e2}$ and $\Delta\boldsymbol{m}$ from six example runs showing high $r(\Delta\boldsymbol{m}, \mathrm{GS})$. The e1 motion regressors were regressed out from the GS and $\Delta\boldsymbol{m}$.

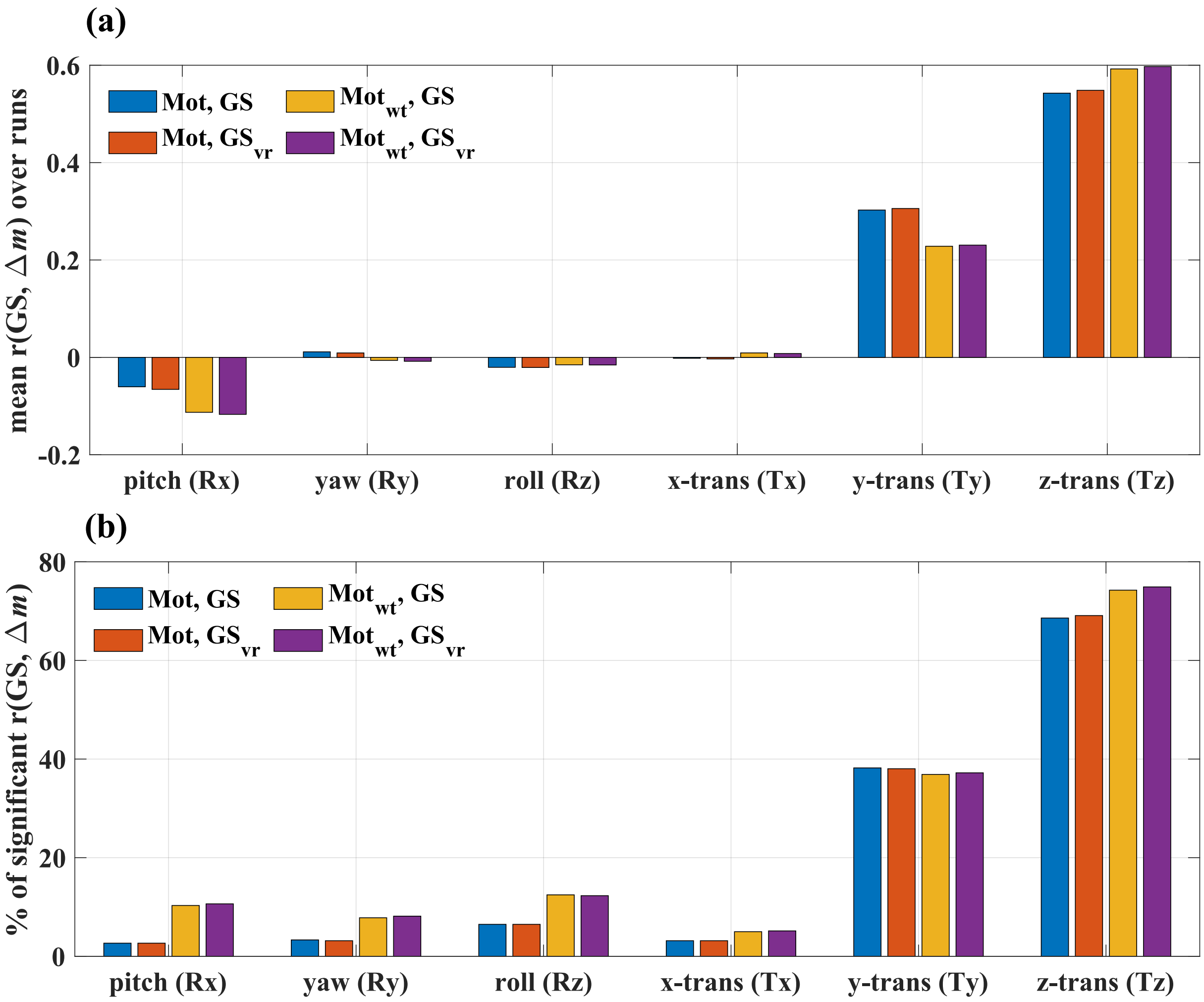


Figure S3: Comparisons of (a) mean $r(\Delta\boldsymbol{m}, \mathrm{GS})$ over runs and (b) the percentage of runs showing significant $r(\Delta\boldsymbol{m}, \mathrm{GS})$ values calculated with combinations of the motion parameters estimated with and without weightings (denoted as Mot and $\mathrm{Mot}_{\mathrm{wt}}$, respectively) and the GS calculated before and after volume registration (denoted as GS and $\mathrm{GS}_{\mathrm{vr}}$, respectively). The $r(\Delta\boldsymbol{m}, \mathrm{GS})$ values were calculated after e1 motion regression. For each motion axis and combination of the GS and motion estimates, an empirical null distribution of $r(\Delta\boldsymbol{m}, \mathrm{GS})$ was generated by the permutation-based method and used to assess the significance of the measured $r(\Delta\boldsymbol{m}, \mathrm{GS})$ values. A Bonferroni-corrected run-wise p-value threshold of 0.05/602 was used, where 602 is the number of runs.

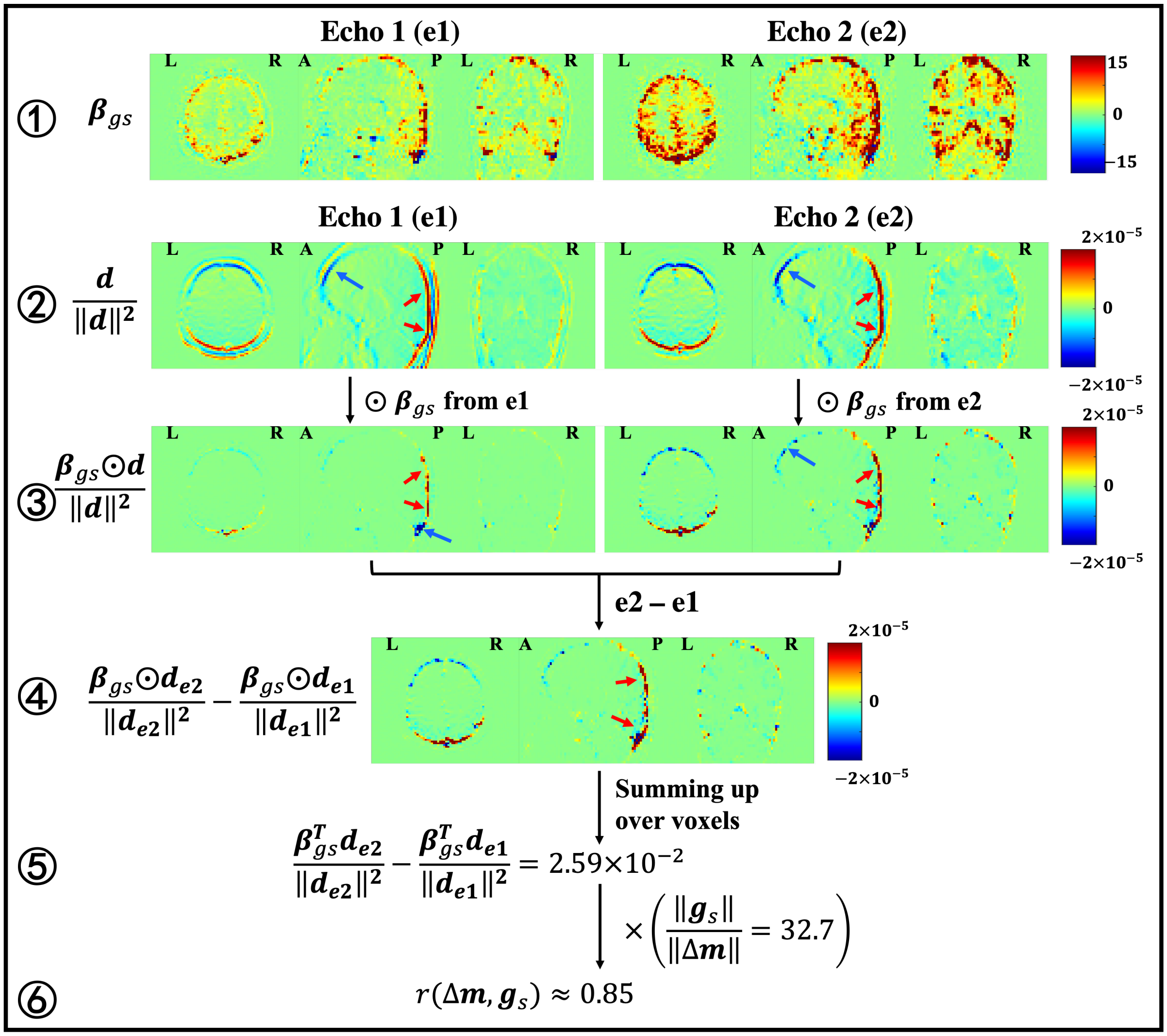


Figure S4: Visualization of the spatial maps underlying $r(\Delta\boldsymbol{m}, \mathrm{GS})$ in the Ty axes from an example run. For each map, three representative slices (one axial, one sagittal and one coronal) are plotted. From top to bottom, rows 1 through 3 show the $\boldsymbol{\beta}_{\boldsymbol{g}_s}$, $\frac{\boldsymbol{d}}{\|\boldsymbol{d}\|^2}$, and $\frac{\boldsymbol{\beta}_{\boldsymbol{g}_s}\odot\boldsymbol{d}}{\|\boldsymbol{d}\|^2}$ maps from e1 and e2, where $\odot$ represents element-wise multiplication. Row 4 shows $\frac{\boldsymbol{\beta}_{\boldsymbol{g}_s}\odot\boldsymbol{d}}{\|\boldsymbol{d}\|^2}$ difference (e2-e1) maps. The red and blue arrows point to the brain regions showing high positive and negative values in the maps, respectively.

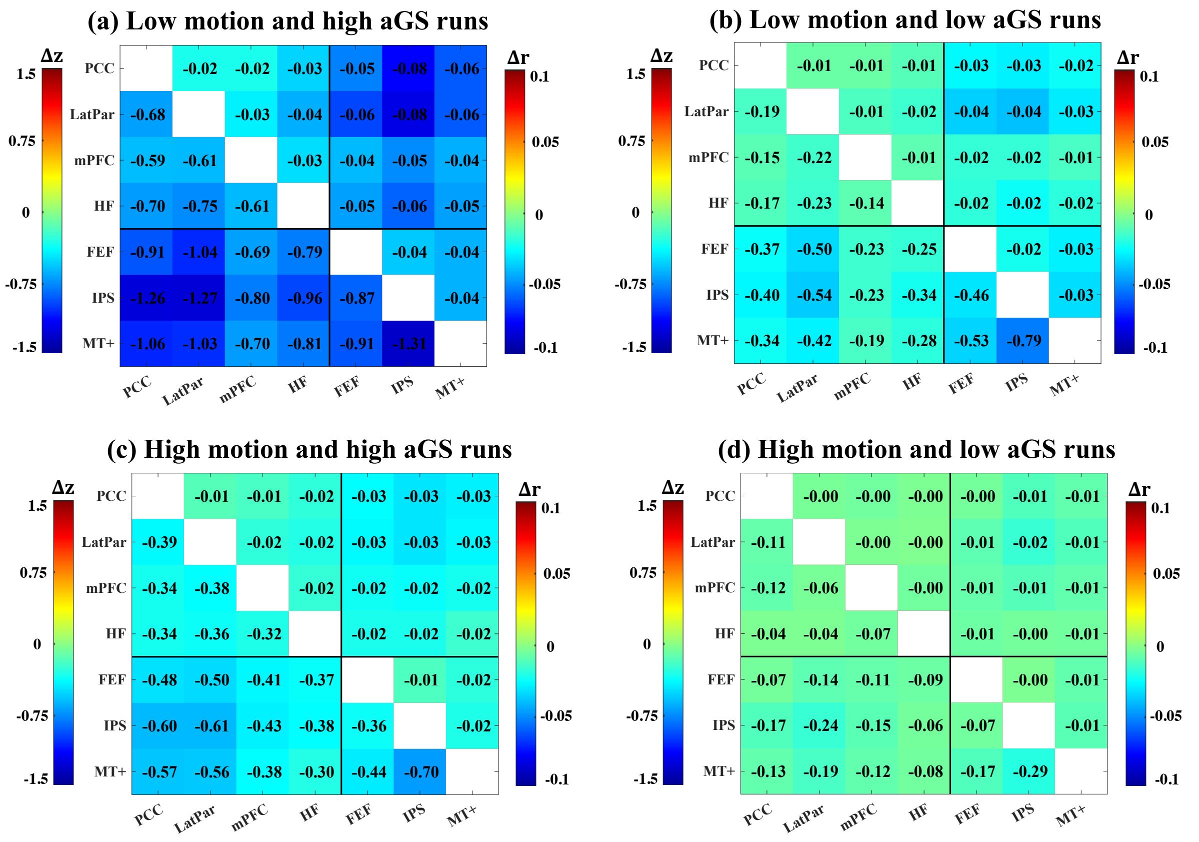


Figure S5: Differences (e2-e1) in ROI-ROI FC calculated after regressing out e1 and e2 motion regressors without motion censoring. The differences were averaged over runs in each of four groups: (a) low motion and high aGS runs, (b) low motion and low aGS runs, (c) high motion and high aGS runs and (d) high motion and low aGS runs. Each subplot is divided into an upper right triangle showing the average differences in r-values and a lower left triangle showing the average differences in z-scores.

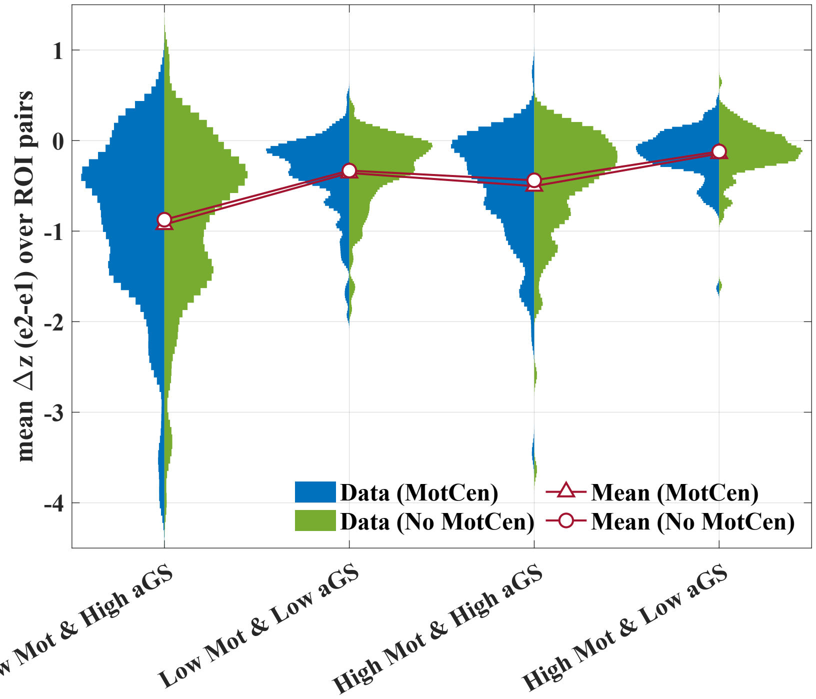


Figure S6: Two-sided violin plots showing the distributions of mean $\Delta z$ (e2-e1) over ROI pairs calculated with (blue) and without (green) motion censoring for (1) low motion and high aGS runs, (2) low motion and low aGS runs, (3) high motion and high aGS runs, and (4) high motion and low aGS runs, where $\Delta z$ indicates the differences (e2-e1) in ROI-ROI FC calculated after regressing out e1 and e2 motion regressors from the e2 data prior to computation of FC. The triangles and circles represent the mean of the distributions. For each group, a paired t-test was calculated to compare the mean $\Delta z$ values with and without motion censoring. The mean $\Delta z$ without motion censoring is significantly ($p < 1 \times 10^{-3}$) less negative than the mean $\Delta z$ with motion censoring for the low motion and high aGS group and the low motion and low aGS group. The effect sizes of the differences are -0.29 and -0.30 for the low motion and high aGS group and the low motion and low aGS group, respectively. No significant ($p > 0.01$) differences were found in the other groups.

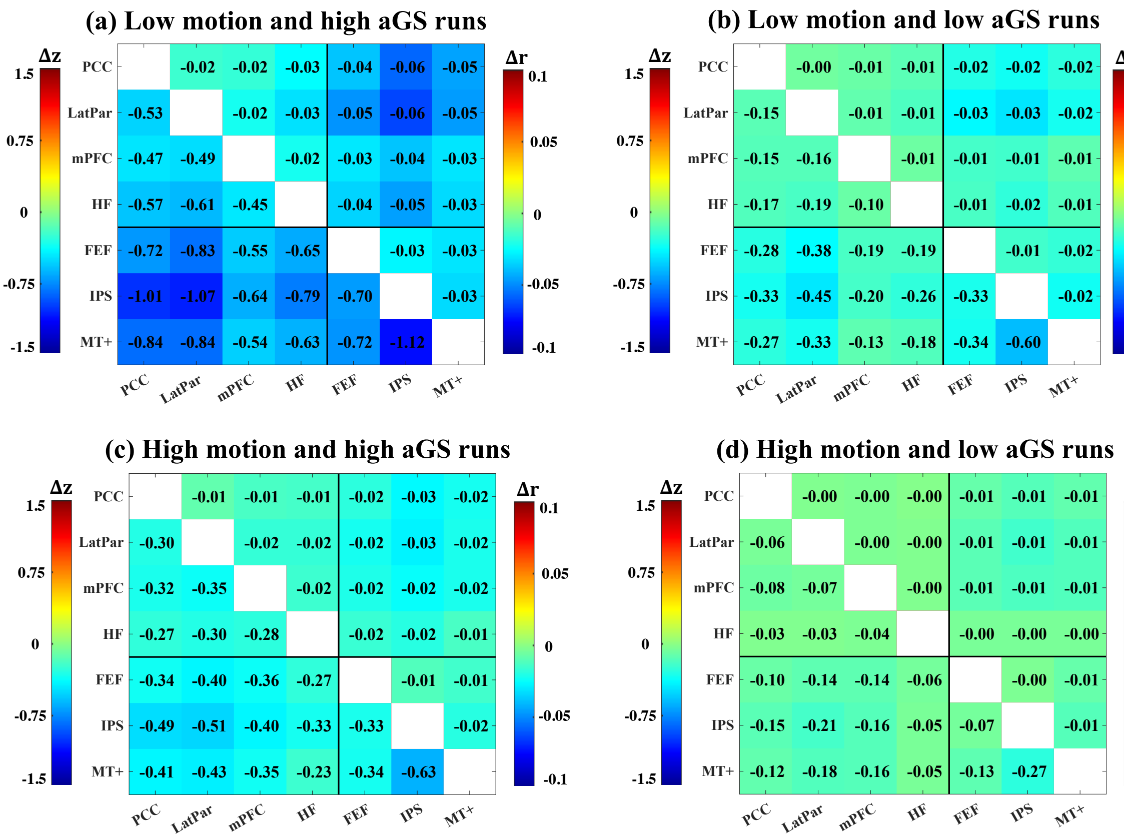


Figure S7: Differences (e2-e1) in ROI-ROI FC calculated after regressing out e1 and e2 motion regressors when including only Ty and Tz motion regressors. The differences were averaged over runs in each of four groups: (a) low motion and high aGS runs, (b) low motion and low aGS runs, (c) high motion and high aGS runs and (d) high motion and low aGS runs. Each subplot is divided into an upper right triangle showing the average differences in r-values and a lower left triangle showing the average differences in z-scores.

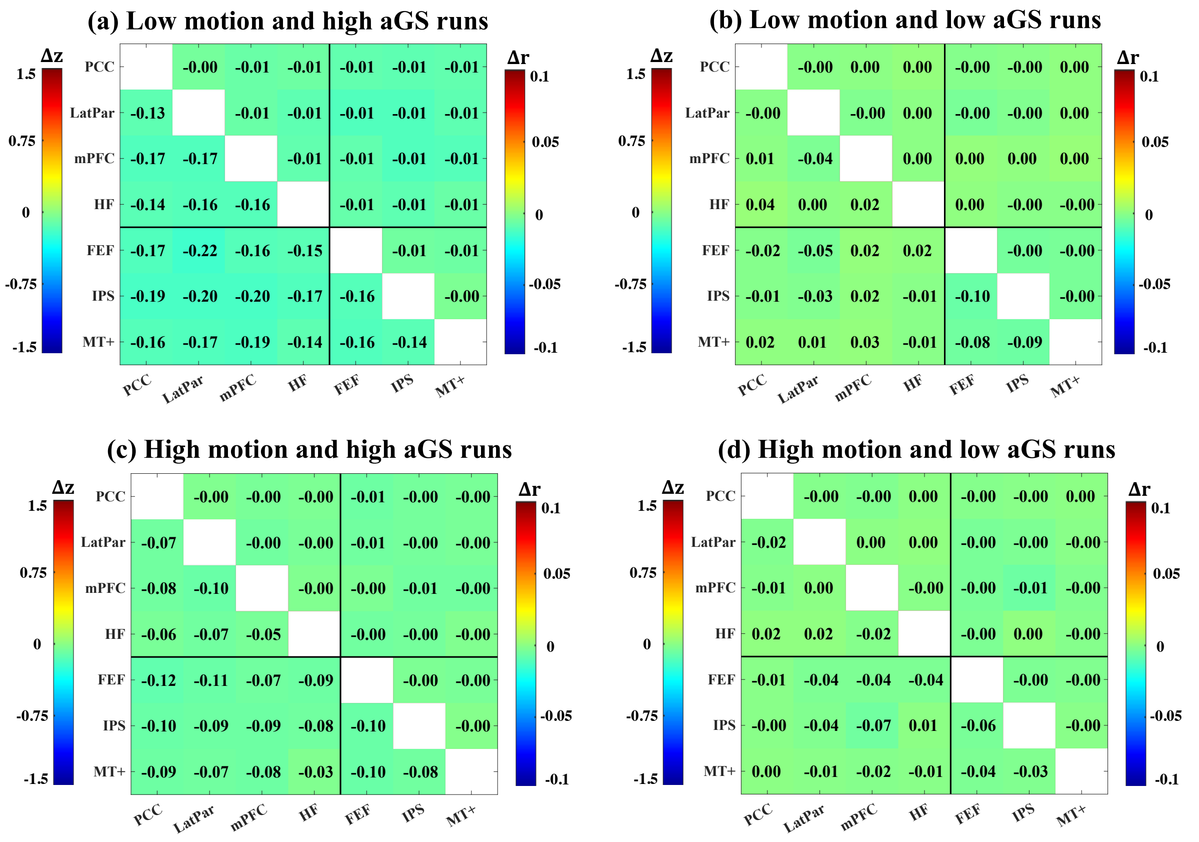


Figure S8: Differences (e2-e1) in ROI-ROI FC calculated after regressing out e1 and e2 motion regressors when excluding Ty and Tz motion regressors. The differences were averaged over runs in each of four groups: (a) low motion and high aGS runs, (b) low motion and low aGS runs, (c) high motion and high aGS runs and (d) high motion and low aGS runs. Each subplot is divided into an upper right triangle showing the average differences in r-values and a lower left triangle showing the averaged differences in z-scores.

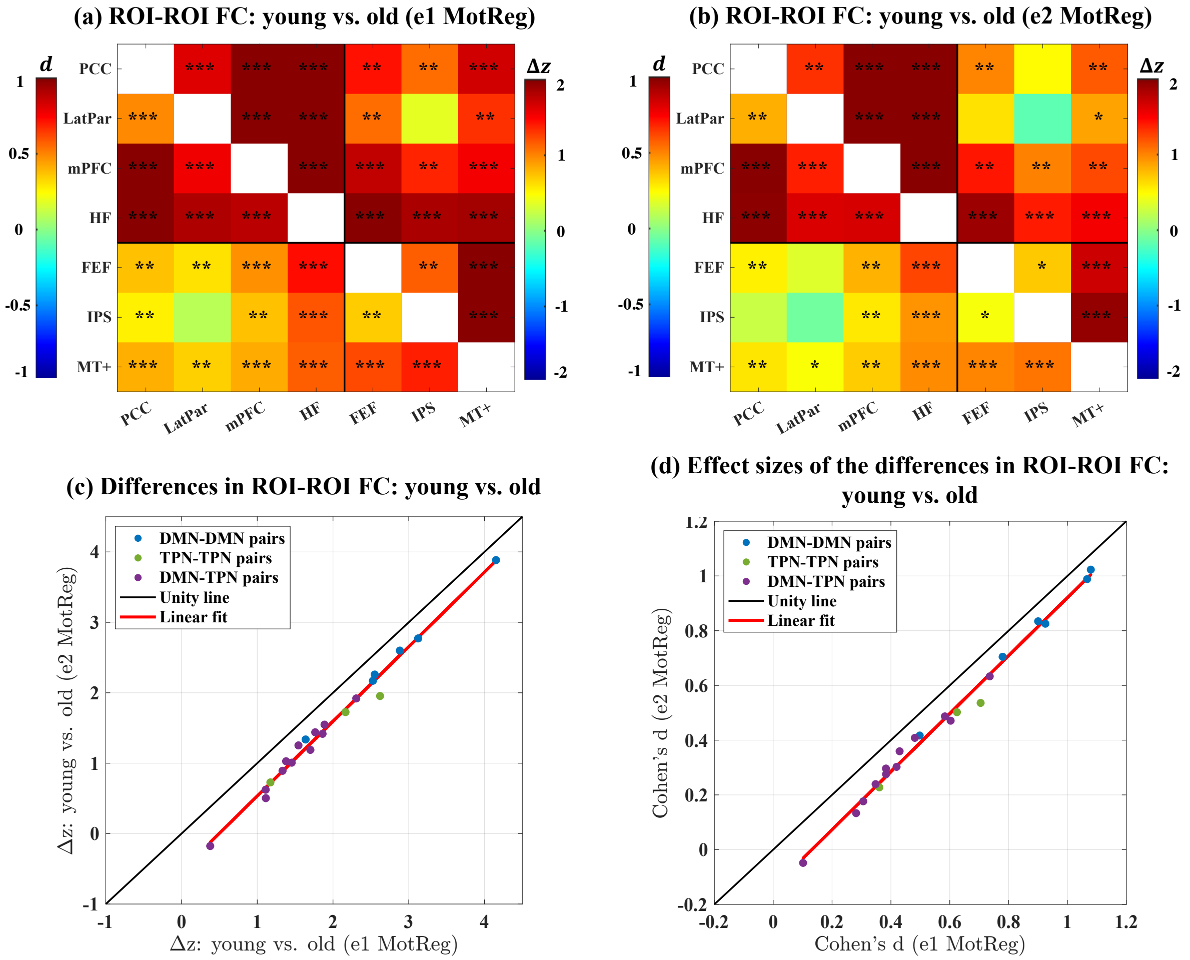


Figure S9: The effect of motion regression on the young vs. old FC differences (young-old) when including only the Ty and Tz motion estimates. (a-b) ROI-ROI FC differences between young and old subjects calculated after (a) e1 and (b) e2 motion regression. Each subplot is divided into an upper right triangle showing the average differences in z-scores between young and old subjects and a lower left triangle showing the effect size of the differences. The black asterisks indicate the statistical significance of the differences assessed by permutation tests (*: $p < 0.01$, **: $p < 1 \times 10^{-3}$, ***: $p < 1 \times 10^{-6}$). A positive value (red color) indicates that the young subjects show higher connectivity as compared to the old subjects. (c) Scatter plot comparing the differences in ROI-ROI FC between the young and the old subjects calculated after e1 and e2 motion regression for all pairs of ROIs. (d) Scatter plot comparing the effect size of the differences in ROI-ROI FC between the young and the old subjects calculated after e1 and e2 motion regression for all pairs of ROIs.

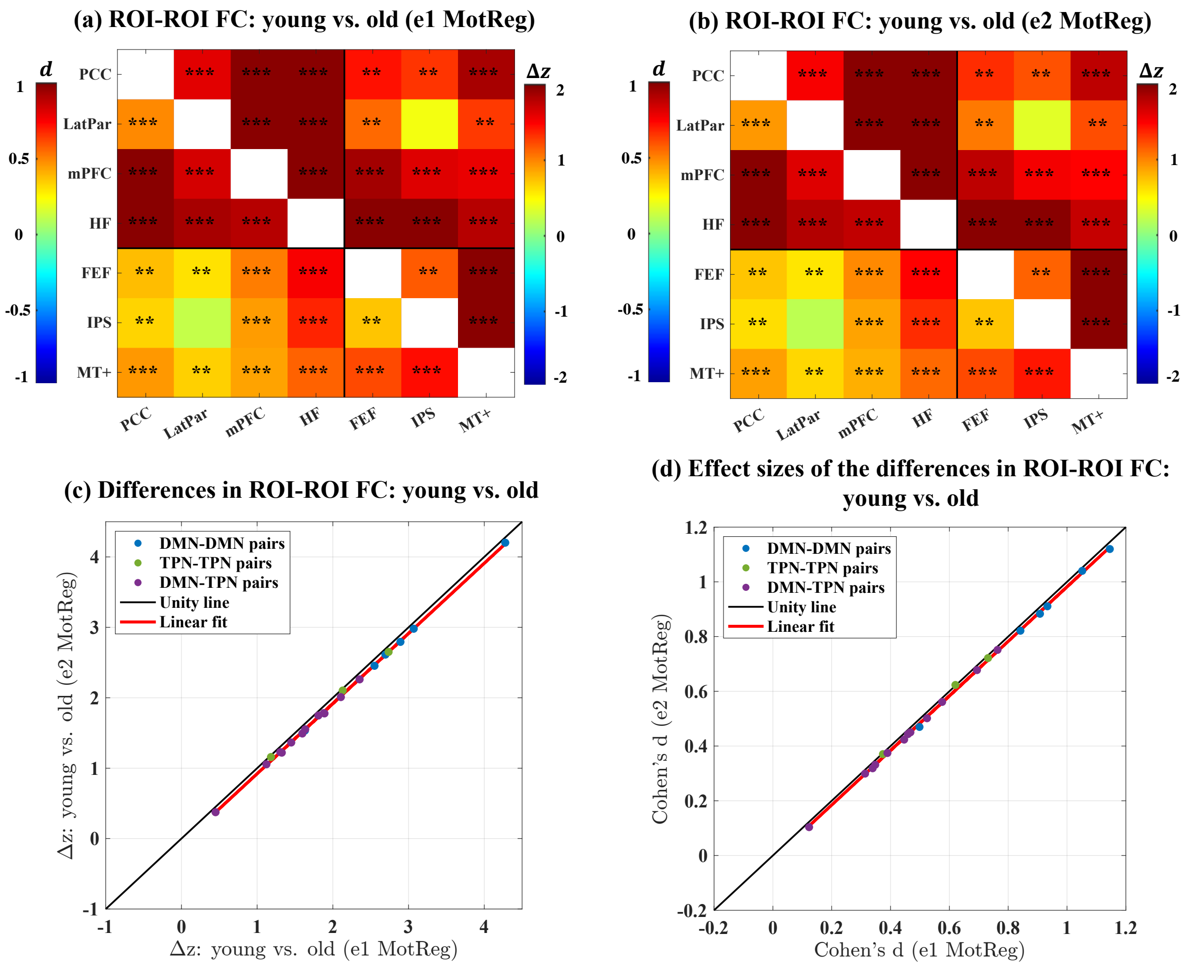


Figure S10: The effect of motion regression on the young vs. old FC differences (young-old) when excluding the Ty and Tz motion estimates. (a-b) ROI-ROI FC differences between young and old subjects calculated after (a) e1 and (b) e2 motion regression. Each subplot is divided into an upper right triangle showing the average differences in z-scores between young and old subjects and a lower left triangle showing the effect size of the differences. The black asterisks indicate the statistical significance of the differences assessed by permutation tests (*: $p < 0.01$, **: $p < 1 \times 10^{-3}$, ***: $p < 1 \times 10^{-6}$). A positive value (red color) indicates that the young subjects show higher connectivity as compared to the old subjects. (c) Scatter plot comparing the differences in ROI-ROI FC between the young and the old subjects calculated after e1 and e2 motion regression for all pairs of ROIs. (d) Scatter plot comparing the effect size of the differences in ROI-ROI FC between the young and the old subjects calculated after e1 and e2 motion regression for all pairs of ROIs.

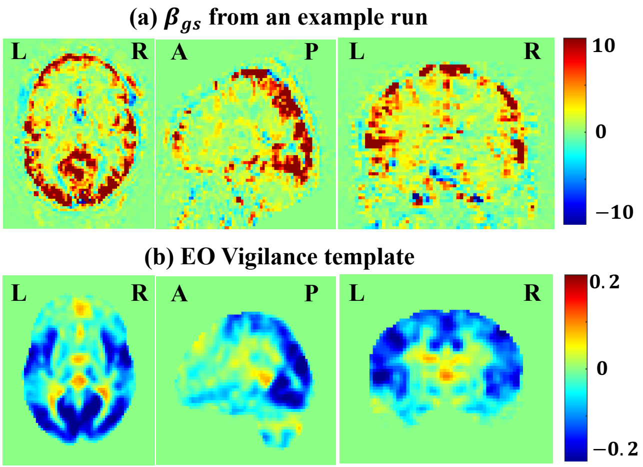


Figure S11: Visualization of (a) $\boldsymbol{\beta}_{\boldsymbol{g}_s}$ from the example run shown in Fig. 3 and (b) the eyes-open vigilance template estimated in (Falahpour et al., 2018). $\boldsymbol{\beta}_{\boldsymbol{g}_s}$ from the example run was calculated with the functional data after registration and transferring to MNI standard space. There is a strong negative correlation ($r = -0.38$, $p < 1 \times 10^{-6}$) between $\boldsymbol{\beta}_{\boldsymbol{g}_s}$ and the vigilance template.

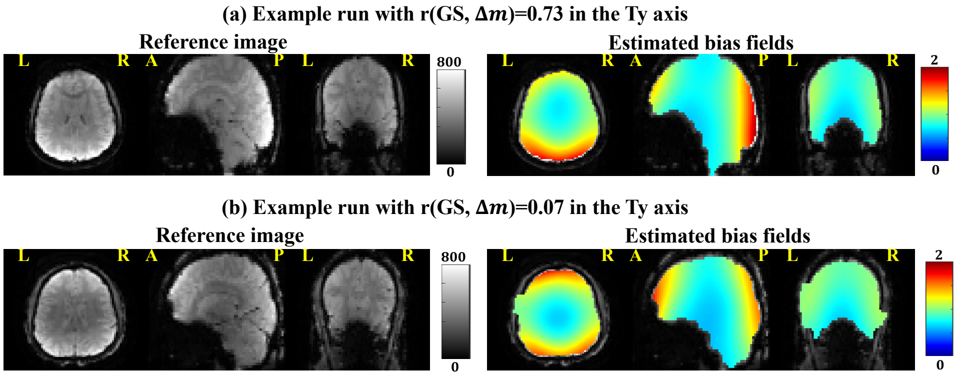


Figure S12: Visualization of the reference images and estimated bias fields from example runs with (a) high $r(\Delta\boldsymbol{m}, \mathrm{GS})$ and (b) low $r(\Delta\boldsymbol{m}, \mathrm{GS})$ in the Ty axis. The bias fields were estimated by SPM segmentation tool and were plotted within brain masks. Both the example runs are classified as low motion and high aGS. For the example run in (a), aGS = 0.61 and mean FD = 0.07mm. For the example run in (b), aGS = 0.80 and mean FD = 0.05mm.

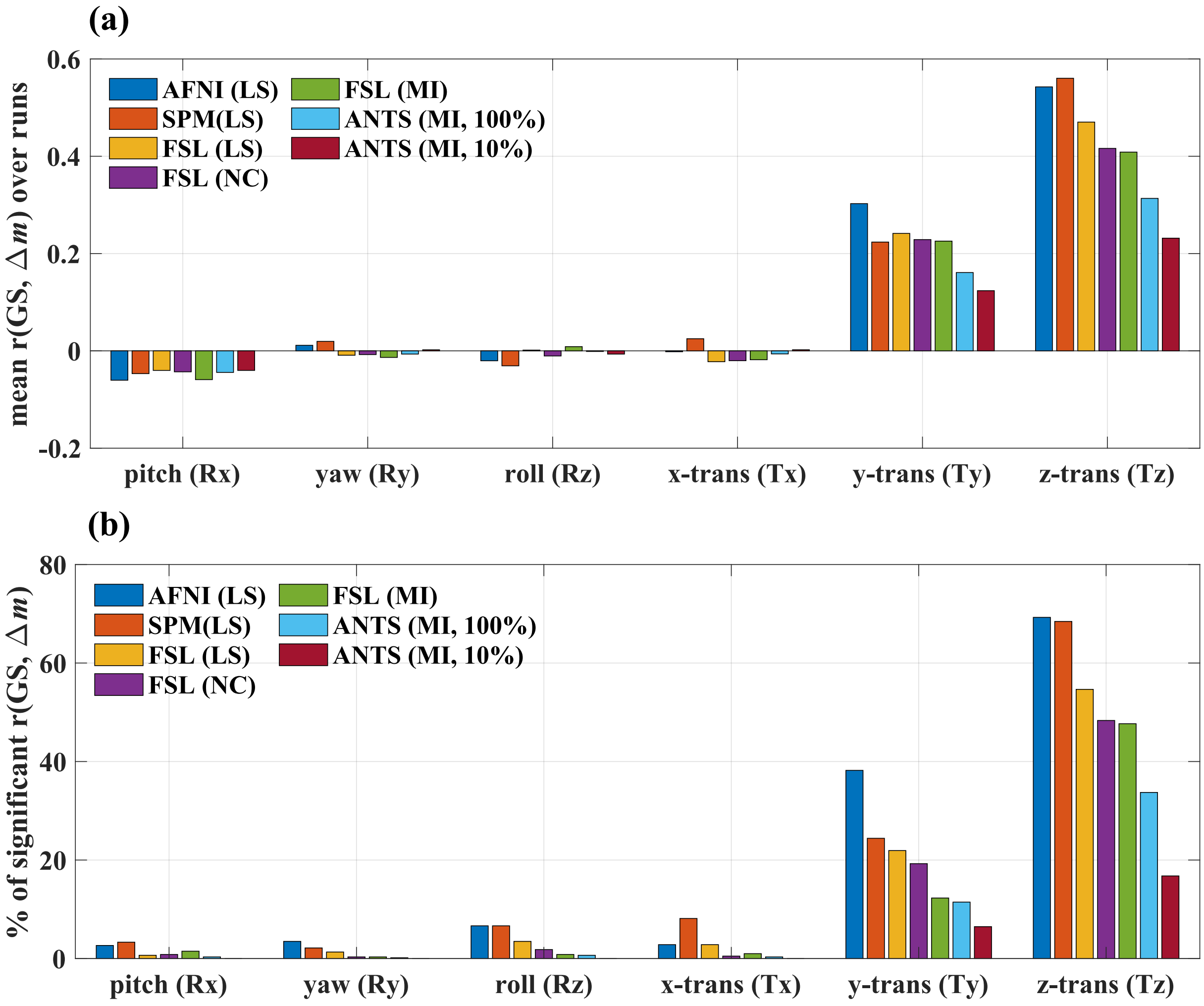


Figure S13: Comparisons of (a) mean $r(\Delta\boldsymbol{m}, \mathrm{GS})$ over runs and (b) the percentage of runs showing significant $r(\Delta\boldsymbol{m}, \mathrm{GS})$ values calculated with motion estimates from different algorithms (AFNI *3dvolreg*, SPM *spm_realign*, FSL *mcflirt*, ANTS *antsMotionCorr*) and cost functions (least-squares (LS), normalized correlation (NC), mutual information (MI)). For ANTS, the sampling percentage was set to either 10% or 100%. The $r(\Delta\boldsymbol{m}, \mathrm{GS})$ values were calculated after e1 motion regression. For each algorithm, cost function and motion axis, an empirical null distribution of $r(\Delta\boldsymbol{m}, \mathrm{GS})$ was generated by the permutation-based method and used to assess the significance of the measured $r(\Delta\boldsymbol{m}, \mathrm{GS})$ values. A Bonferroni-corrected run-wise p-value threshold of 0.05/602 was used, where 602 is the number of runs.

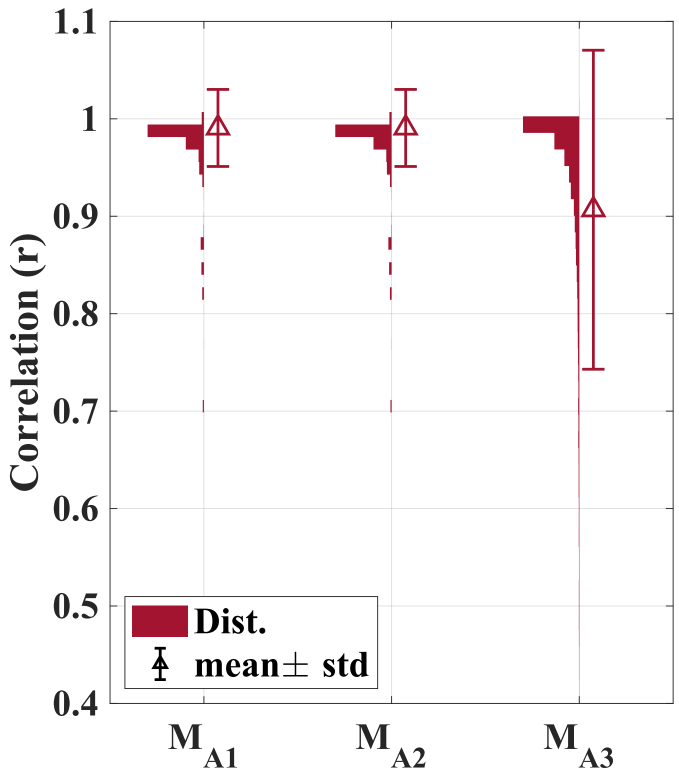


Figure S14: Distributions of the correlations between the motion estimates from AFNI *3dvolreg* and the approximated motion estimates, i.e. $\boldsymbol{M}_{Ai}$, $i = 1, 2$ and 3. The approximations are described in Eq. B.1. The distributions include the correlations for all six motion axes from all runs for both e1 and e2.

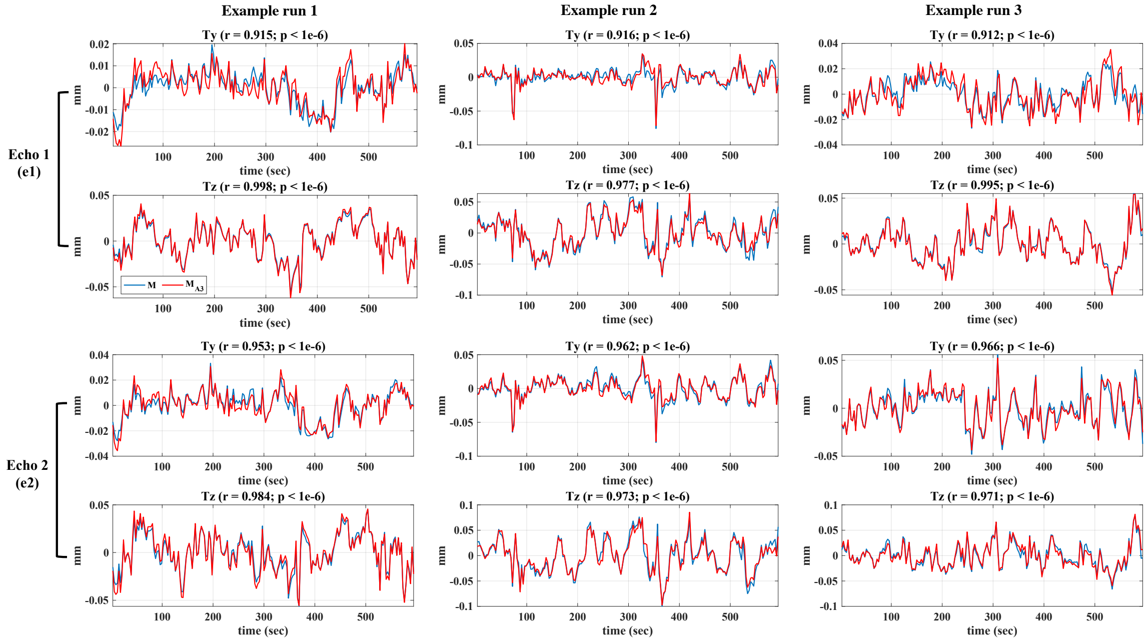


Figure S15: Comparison of the motion estimates from AFNI *3dvolreg* (i.e. $\boldsymbol{M}$, blue) and the motion estimates after the third approximation (i.e. $\boldsymbol{M}_{A3}$, red) in the Ty and Tz axes from three example runs. The top two rows show the motion estimates for e1, and the bottom two rows show the motion estimates for e2.
